# Opsin-based photoreception in Crinoids: a molecular and behavioural study of *Antedon bifida*

**DOI:** 10.1101/2024.08.14.607903

**Authors:** Youri Nonclercq, Marjorie Lienard, Alexia Lourtie, Emilie Duthoo, Lise Vanespen, Igor Eeckhaut, Patrick Flammang, Jérôme Delroisse

## Abstract

Opsins are essential photoreceptor proteins that enable both visual and nonvisual light perception in metazoans. Although extensively studied in eyed invertebrates, their role in eyeless organisms such as echinoderms remains underexplored. Within the echinoderm phylum, studies have primarily focused on sea stars, sea urchins, and brittle stars, leaving crinoids, the most basal echinoderm lineage, entirely unexplored. Currently, only a limited number of behavioural observations have suggested that crinoids may possess light sensitivity. This study investigated the behavioural, morphofunctional, and molecular foundations of opsin-based photoreception in *Antedon bifida,* a European crinoid species from the comatulid order. In this context, the behavioural response to different light wavelengths, characterisation of opsin genes in the recent chromosome-scale genome of this species, opsin immunolocalisation within the crinoid tissues, and *in vitro* functional characterisation of opsins have been investigated.

*In vivo* tests indicated significant negative phototactic behaviour induced by a wide range of light wavelengths (463–630 nm) with maximum sensitivity to blue light (λ_max_ = 463 nm). *In silico* genome analyses revealed the presence of only three rhabdomeric opsin genes, located on chromosomes 4 (Abif-opsin 4.1) and 6 (Abif-opsin 4.2 and 4.3). All crinoid opsins were phylogenetically clustered as a sister group to all other echinoderm rhabdomeric opsins, supporting their evolution via duplication of an ancestral gene in the crinoid lineage. The low opsin diversity contrasts with that of other echinoderms, which are generally characterised by up to eight bilaterian opsin types. Interestingly, *A. bifida* opsin sequences present typical amino acid residues of rhabdomeric opsins of other bilaterians, including two conserved cysteines (C110 and C187), the probable ancestral E181 counterion, the typical G-protein signalling NPxxY(x)_6_F pattern, a highly conserved lysine potentially covalently bound to a chromophore, and the (D)RY motif, all of which support a photoreceptive function. Heterologous *in vitro* expression in HEK293T cell cultures subsequently confirmed the photoreceptive function of all three *A. bifida* opsins. Indeed, these three crinoid opsins formed active complexes with 11, cis-retinal association, and once purified, showed different absorbance peaks in the short wavelengths of the visible light spectrum ranging from 425 to 520 nm. Finally, immunoreactivity to newly generated antibodies against sea star opsins highlighted two potential crinoid opsins in several tissues associated with the ambulacral grooves of the calyx and pinnules. Within these tissues, one Abif-opsin is potentially expressed both in the ectoneural basiepithelial nerve plexus and in the hyponeural nerve plexus. However, a different opsin is also expressed in the sensory papillae of tube feet. Localisation of at least two opsins in different sensory structures suggests the presence of a complex extraocular photoreception system based exclusively on rhabdomeric opsins in this crinoid species.

## Background

Light plays a significant role in marine ecosystems, and most marine organisms have specific photosensitive organs such as the eyes, which contain opsins, the prototypic light-sensitive GPCR receptors of eumetazoans. Interestingly, echinoderms often lack specialised visual organs except for optic cushions located at the tip of sea star arms [Takasu and Yoshida 1983; Johnsen 1997; Garm and Nilsson 2014; Petie et al. 2016] and putative eyespots in a few holothurian species [Berrill 1966, Yamamoto and Yoshida 1978]. However, numerous behavioural observations have confirmed that most investigated echinoderms are light-sensitive [for review, Yoshida et al. 1984, Sumner-Rooney and Ullrich-Lüter, 2023]. In many species, extraocular photoreception is mediated by different classes of opsins expressed in different tissues intimately connected to the nervous system. Two main opsin types, ciliary and rhabdomeric, have been identified in various adult anatomical structures, such as the tube feet, integument, and spines of sea urchins [Ullrich-Lüter et al. 2011; 2013], sea stars [Delroisse et al. 2013, Ullrich-Lüter et al. 2013, Clarke et al. 2024] and brittle stars [Moore and Cobb, 1985; Cobb and Hendler, 1990; Delroisse et al. 2014; 2016; Sumner-Rooney et al. 2018; 2020; 2021]. In addition, the optic cushion of sea stars specifically expresses several opsin types [Ullrich-Lüter et al. 2011, Lowe et al. 2018]. A third type of opsin, GO-opsin, has also been identified in two clusters of sensory cells on either side of the apical organ of pluteus sea urchin larvae [Valero-Gracia et al. 2016; Valencia et al. 2021; Cocurullo et al. 2023]. Extensive phylogenetic analyses of bilaterian opsins have shown that echinoderms are one of the phyla with the largest diversity of ancestral opsin types, with seven opsin classes on a total of nine predicted in the common ancestor of all bilaterians [D’Aniello et al. 2015; Ramirez et al. 2016]. Some ancestral opsin types, such as bathyopsins and chaopsins, are currently almost exclusively restricted to echinoderms [Ramirez et al. 2016].

The crinoid group has been the least studied echinoderm class in terms of sensory perception in general and photoreception in particular. However, given that Crinoidea currently represents the most basal clade of echinoderms, separated from the other orders since at least the Lower Ordovician period (480 million years ago), their phylogenetic position makes them an interesting group to better understand the evolution of photoreception, not only within the whole phylum. It has long been known that some crinoid species are sensitive to light. Indeed, ethological observations have shown that several species of comatulids (an order of crinoids, most of which are non-pedunculated in the adult stage) living at shallow depths are predominantly nocturnal, extending their arms to filter feed at night [Magnus 1964; Rutman and Fishelson 1969; Meyer 1973; Vail 1987]. However, when these organisms are exposed to daylight, they tend to hide in reef crevices or under shaded underwater overhangs. Other observations point to the rapid retraction of the arms when exposed to strong sunlight [Meyer 1973]. However, this nocturnal activity is less systematically observed in individuals living at greater depths (beyond 10-20 m below the surface), where the luminosity of the environment is lower [Fishelson 1974]. Some studies have also reported swimming escape behaviour in feather stars, such as the European species *Antedon bifida,* when exposed to strong light stimuli [Dimelow 1958]. However, these isolated observations leave many unexplored areas, such as a deeper characterisation of the photoreceptive behaviours or a determination of the spectral sensitivity of these animals. No photosensitive sensory structure has been formally demonstrated in extant crinoids. The repertoire of opsin genes remains mostly unexplored, with only one rhabdomeric-type opsin identified in two transcriptome datasets from *Antedon mediterranea* and *Florometra serratissima* [D’Aniello et al. 2015].

The highly specialised lifestyle of crinoids and their phylogenetic position make them a prime group for better understanding the evolution of the large opsin diversity in echinoderms, as well as the evolution of their extraocular photoreception.

This study provides new insights into the mechanisms of opsin-based photoreception in crinoids by focusing on the phototactic behavior of the feather star species *Antedon bifida*. This small, widespread species inhabits shallow waters along the North Atlantic European coasts, ranging from just below the surface to depths of 200 meters. It is typically found during daylight hours in shaded, rocky environments, avoiding full light exposure. By leveraging a recently available chromosome-scale genome, we annotated and described the complete repertoire of opsin genes in *A. bifida*, enabling the in vitro expression of these photoreceptor proteins and the determination of their light absorption spectra. Additionally, the precise anatomical localisation of several opsin-based photoreceptors allowed us to characterise the photosensory structures responsible for extraocular photoreception in this species. Behavioral tests revealed that *A. bifida* exhibits significant negative phototactic behavior, particularly in response to blue light. This response is likely mediated by three rhabdomeric opsins, which demonstrate functional photosensitivity to short wavelengths. Our combined approaches uncovered novel candidate photosensitive tissues and illuminated the molecular underpinnings of light perception in this understudied echinoderm class. These findings not only enhance our understanding of crinoid photoreception but also pave the way for future molecular and functional studies, offering broader insights into the evolutionary adaptations of light perception within the echinoderm phylum.

## Material and Methods

### Collection of crinoid specimens

Antedon bifida specimens **(Fig. 1A)** were collected from two harbours on the Crozon peninsula, Brittany, France **(Fig. 1B-D)**. The organisms were abundant, clinging to the undersides of mooring pontoons just below the water’s surface **(Fig. 1E, F)**. Approximately 30 individuals were brought back to the University of Mons (Belgium) and kept in an aquarium containing approximately 400 L of seawater at 15°C (with a salinity of 34 PSU) in a closed circuit, following a 12H/12H day/night cycle **(Fig. 1G)**.

**Figure 1.**
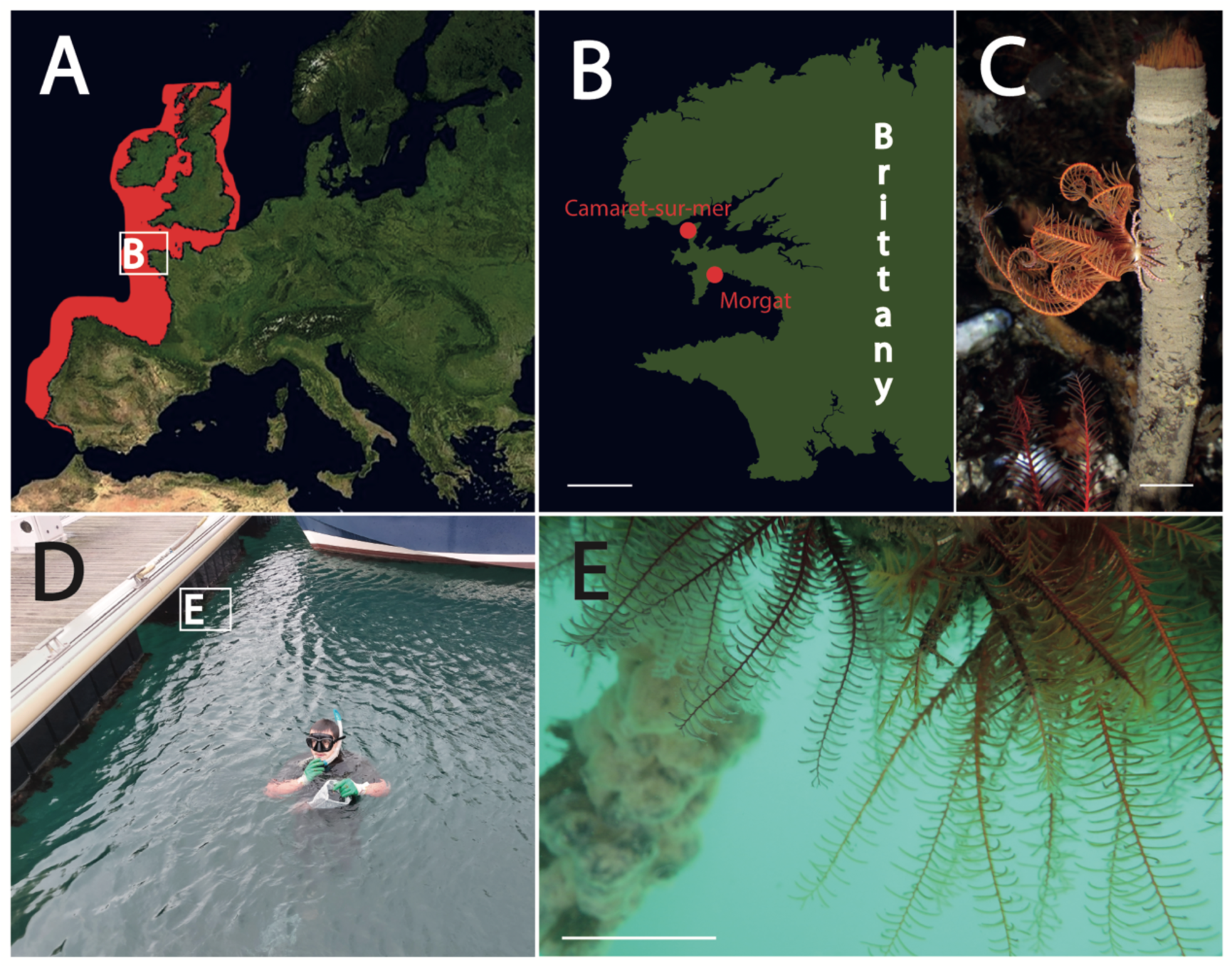
Distribution, habitat, and collection sites of *Antedon bifida*. (A) Geographical distribution of *A. bifida* along the North Atlantic coast of Europe. (B) Focus on two collection sites (Morgat and Camaret-sur-Mer) located on the Crozon Peninsula in Brittany (France). (C) Adult specimen of *Antedon bifida* clinging to a sabellid tubeworm. (D) Collection site in the marina of the Camaret-Sur-Mer. (E) Dense population of *A. bifida* hanging under mooring pontoons. Scales: B. 15km, C. 20mm, E. 20mm.

### Ethological experiments in *Antedon bifida*

Phototactic behaviour tests were carried out using an elongated experimental aquarium (50 cm long and 7.5 cm wide) with three sides rendered opaque **(Fig. 2)**. A monochromatic light source (LED bulb GU10 of 1 Watt) was placed against the transparent glass end of the experimental device to create a light gradient across the entire aquarium. The light intensity along this gradient was measured with a light metre (Digital Lux Meter AR823^+^, Smart Sensor) and the brightness values varied from 46,000 Lux (6.7 µW/cm^2^) next to the light source to 3,600 Lux (0.5 µW/cm^2^) at the opposite extremity. The mean value in the middle part of the aquarium was 10,000 Lux (1.5 µW/cm^2^). The floor was covered with a rigid plastic mesh, allowing the crinoids to cling and move easily in the aquarium. Three different light wavelengths were tested: blue light (λ_max_= 463 nm, half-peak bandwidth HBW= 27 nm), green light (λ_max_= 512 nm, HBW= 33 nm), and red light (λ_max_= 630 nm, HBW= 20 nm). The spectra of each LED lamp were precisely measured using a spectrometer (Ocean View, FLAM-S-UV-VIS spectrometer). One test was carried out with polychromatic white light, and a negative control was used without a light source. Each of these five experimental conditions was carried out on 10 individuals of *A. bifida* during the same time of the day in a room completely immersed in the dark. Individuals were tested separately to avoid aggregation. Each crinoid specimen was first conditioned in the dark for approximately 5 min. Then, for each test, individual was placed in the centre of the aquarium (25 cm from the illuminated end) and its displacement towards (positive displacement) or against (negative displacement) the light source was measured after 10 min. These displacement measurements were based on the position of the crinoid calyx centre relative to its initial position. Statistical tests were used to confirm the homogeneity variance and normal distribution of the displacement data using Bartlett and Quantil/Quantil tests, respectively. The mean values of the distance covered for each light wavelength tested were then compared to the mean distance travelled by individuals without illumination (negative control) using a one-way analysis of variance (ANOVA), followed by a Dunnett post hoc test with RStudio statistical software.

**Figure 2.**
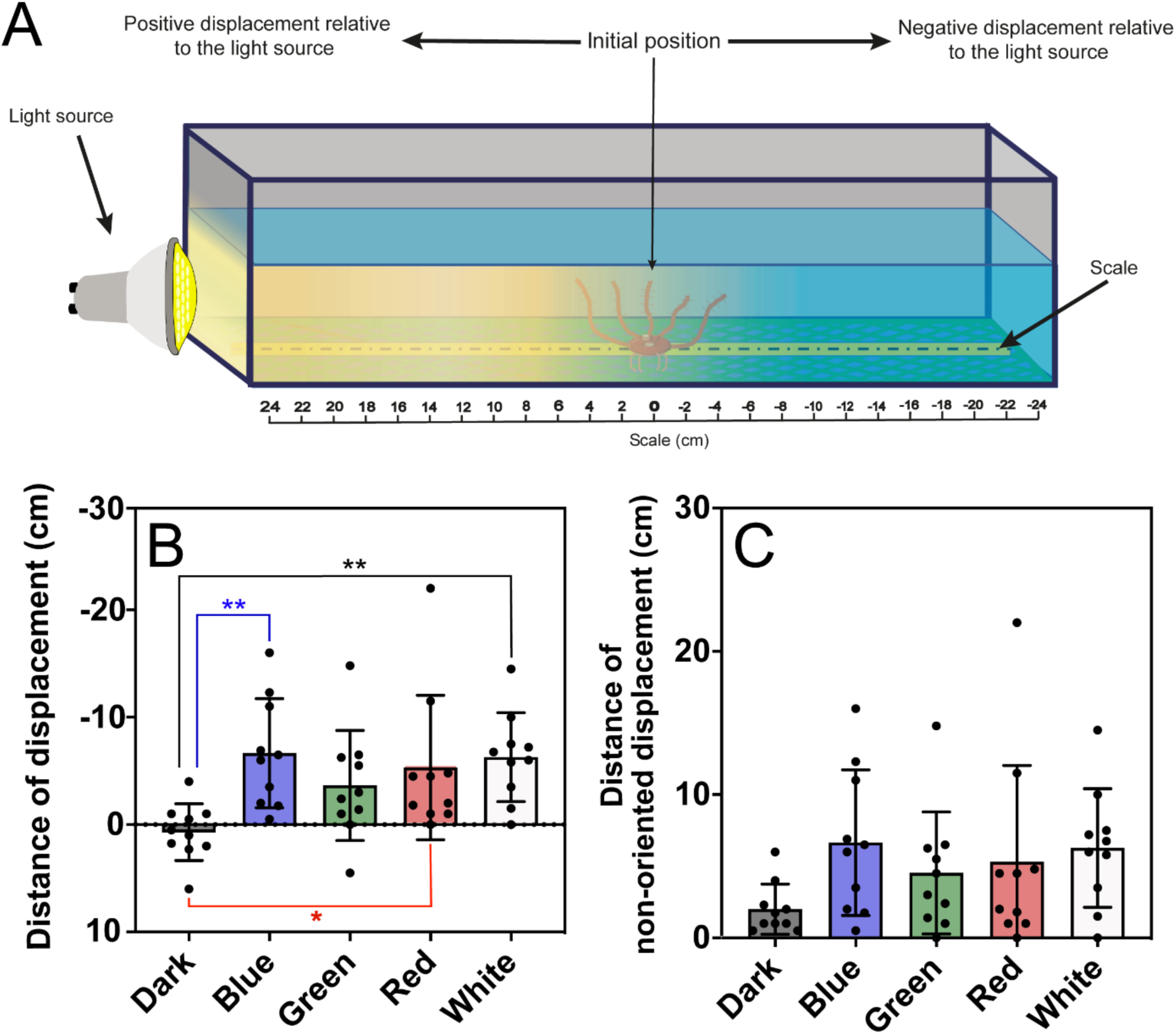
Phototactic behaviour in *Antedon bifida*: (A) Schematic representation of the experimental aquarium with a light gradient. (B) Graph showing the distance travelled by individuals within the experimental set-up over 10 minutes of time following different light stimuli (Black = no light; Blue = λmax of 463 nm; Green = λmax of 512 nm; Red = λmax of 630 nm and White = polychromatic white light). Negative displacement indicated a displacement away from the light source. One-factor ANOVA and Dunnett’s post hoc statistical tests showed that crinoids exhibited a significant oriented displacement towards the darker region of the light gradient for white, blue and red light compared with the no-light control. (C) Graph showing the distances covered by the individuals in absolute values, showing the rate of movements of crinoids depending on the type of light stimulus. Dunnett’s one-factor ANOVA and post hoc tests did not provide significant results between no-light controls versus different wavelengths tested.

### *In silico* identification of opsin genes in *crinoids*

The diversity of the opsin genes in our model crinoid species, *A. bifida*, was determined using a homology-based approach. This method uses several published opsin reference sequences as templates [D’Aniello et al. 2015, Ramirez et al. 2016], *i.e.* opsins from the echinoderm species, the sea urchin *Strongylocentrotus purpuratus,* and opsins from additional representative bilaterians **(*Suppl. Table S1*)**.

Homology "tBLASTn" searches of the predicted protein sequences were performed against the assembled *A. bifida* reference genome (ecAntBifi1.1) available in the NCBI database. The following tBLASTn parameters were used: *Matrix: Blosum62; gap costs: existence 11, extension 1*. We then applied a cut-off e-value < 1^e-05^ for putative opsin candidates showing a strong similarity to two highly conserved regions in functional opsins: the 7^th^ transmembrane helix corresponding to the Schiff base binding zone of retinal (equivalent K296 lysine residue in the bovine rhodopsin), and the G-protein binding zone. Since the genome of *A. bifida* has not yet been annotated, a reciprocal BLAST of these selected sequences was then performed on the entire NCBI database to validate their opsin identification.

To obtain the most complete and accurate opsin gene predictions, we used several gene prediction softwares like GENSCAN [Burge and Karlin 1997: http://hollywood.mit.edu/GENSCAN.html], AUGUSTUS [Oliver et al. 2011: https://bioinf.uni-greifswald.de/augustus/] and FGENESH [Solovyev et al. 2006: http://www.softberry.com/berry.phtml?topic=index&group=programs&subgroup=gfind] using tBLASTn on all chromosome portions of the *A. bifida* genome identified to contain opsin loci.

Then, we carried out a manual curation as follows: for each predicted opsin, only the longest portion of the protein sequence which, using a MAFFT alignment (MAFFT version 7, Research Institute for Microbial Diseases, Japan, https://mafft.cbrc.jp/alignment/server/index.html) [Katoh et al. 2019] and maximising the high similarity to other echinoderm opsin reference sequences **(*Suppl. Table S1*)** has been selected as the most accurate crinoid opsin sequence.

Next, a sequence similarity comparison between opsin sequences found in *Antedon bifida* and the reference opsin repertoire of the sea urchin *S. purpuratus* (Sp-opsin 1; 2; 3.1; 3.2; 4; 5; 6; 7 and 8) was performed using SIAS webtool (Sequence Identity And Similarity, Universidad Complutense de Madrid) (http://imed.med.ucm.es/Tools/sias.html).

The chromosomal locations of *A. bifida* opsin genes within the genome and the number of intron/exon regions were finally determined using internal tBLASTn search of the candidate protein sequences identified above against the *A. bifida* genome. Gene positions within chromosomes were visualised using IGV (Integrative Genomics Viewer) genome browser [Robinson et al 2023]. The number and position of introns and exons were annotated by combining tBlastn, FGENESH in Softberry [http://www.softberry.com], and the Kablammo software [Wintersinger and Wasmut 2014, https://kablammo.wasmuthlab.org].

A similar procedure was carried out to determine all opsin sequences of two additional crinoid species: *Anneissia japonica* (Comatulidae) and *Nesometra sesokonis* (Antedonidae like *A. bifida*), whose reference genomes are also present in the NCBI genomic database (ASM1163010v1 and ASM2563120v1, respectively).

### Phylogenetic analyses of crinoid opsins

A phylogenetic analysis was carried out on opsin protein sequences previously discovered in the two crinoid model species (*A. bifida*, *N. sesokonis*) to highlight their relationship with other opsins. For this analysis, 111 different protein sequences from 21 species representing the 5 echinoderm classes were used alongside opsin sequences of a few other representative groups of bilaterians. These sequences were selected based on public metadata [Delroisse et al. 2014; D’Aniello et al. 2015; Ramirez et al. 2016; Lowe et al. 2018] and genomic databases such as NCBI to represent all nine ancestral opsin lineages present in bilaterians (ciliary opsin, bathyopsin, Go-opsin, canonical and non-canonical Rhabdomeric opsin, chaopsin, peropsin/RGR opsin, neuropsin and xenopsin) **(Suppl. Table S1)**. Eight additional melatonin receptor protein sequences were added as outgroups. The sequences were then aligned using MAFFT version 7 [Katoh et al. 2019]. The alignment was then trimmed using the TrimAl software tool hosted on the NGPhylogeny.fr online platform (https://ngphylogeny.fr/tools/tool/284/form). The tree was constructed using IQ-Tree (http://iqtree.cibiv.univie.ac.at/) [Nguyen et al. 2015] based on maximum likelihood analysis with 1000 ultrafast bootstraps. The best-fit model recommended by ModelFinder [Kalyaanamoorthy et al. 2017] to our unpartitioned dataset was LG+F+I+G4, and the number of parsimony informative sites was 726 on the 1583 amino-acid sites used in total. Tree visualisation was finally performed on the iTOL Interactive Tree Of Life platform (https://itol.embl.de/).

### *In silico* modelling of crinoid opsins

*A. bifida* opsin protein sequences were uploaded on Swiss model (Waterhouse et al 2018) to generate homology structures using the squid rhodopsin 2ziy before visualisation and analysis in PyMOL.

### Opsin heterologous expression and functional characterisation

#### In vitro expression in HEK293T cells

Opsin open reading frames were synthesised and codon-optimised for mammalian expression by Genscript, subcloned in a custom pcDNA5-FLAG-T2A-mRuby2 expression vector, expressed transiently in HEK293T cells and purified in the presence of 11 *cis*-retinal following the procedure detailed in Liénard et al 2021. Briefly, twenty plates of 4.10^6^ HEK293T cells each were transfected with 24 μg endotoxin-free plasmid construct in a ratio of 1:2.5 DNA:PEI (polyethylenimine 1mg/mL in water) before delivery of 11,*cis-*retinal (5 μM final concentration) under dim light and incubation in the dark for 48 hours. Cells were harvested and pelleted in 50mL cold Hepes buffer (3 mM MgCl_2_, 140 mM NaCl, 50 mM Hepes pH 9.0 or 7.0). The total membrane protein fraction was extracted with gentle rotation at 4°C in 10 mL extraction buffer (3 mM MgCl_2_, 140 mM NaCl, 50 mM Hepes pH 7.0 or 9.0, 20% glycerol vol/vol, 1% n-dodecyl β-D-maltoside, complete EDTA-free protein inhibitors) in presence of 40 μM *cis*-retinal, and the crude extract was centrifuged. Rhodopsin complexes in the supernatant were bound to FLAG resin (Sigma-Aldrich) with gentle rotation overnight at 4°C. Elution was then performed at room temperature following the addition of 0.75 mg 3xFLAG peptide (Sigma-Aldrich), then the eluate was concentrated using an Amicon Ultra-2 centrifugal filter unit with Ultracel-10 membrane (Millipore) for 1 hour at 4,000 rpm at 4°C. Ultraviolet-visible (UV-VIS) absorbance was recorded using an Implen NP80 nanophotometer and analysed in RStudio (V2021.09.2) using custom scripts for polynomial visual template analysis to obtain the dark absorbance spectrum [Liénard et al 2022].

#### Immunoblot analysis

Protein purification fractions were collected and loaded on a 4-15% MP TGX (Biorad). Proteins were separated at 4°C and 80V for 1h40 min, transferred to a nitrocellulose membrane on a TurboBlotTransfer system (Biorad Laboratories) and the membranes were blocked for 1 hour with 5% milk (Biorad laboratories) in tris-buffered saline containing 0.1% Tween 20 (TBS-T, Biorad Laboratories). Membranes were then incubated overnight with monoclonal anti-FLAG M2 antibody 1:2,500 (F1804, Sigma-Aldrich) on a gently rocking platform at 4°C. The membranes were washed in TBS-T, incubated with HRP Conjugated ECL anti-mouse (1:1000; NA931, Sigma-Aldrich) for 1 hour at room temperature, washed in TBS-T, revealed using the SuperSignal West Femto (Thermo Scientific) and imaged on a ImagQuant800 (Cytiva Amerhsam).

### Histology

#### Specimen preparation

*Antedon bifida* specimens intended for histological and immunohistochemical analysis were anaesthetised with a 3.5% magnesium chloride in sea water solution for 2 min before being placed 4 h in a non-acetic Bouin fixative fluid (75% of a picric acid saturated solution and 25% of formaldehyde commercial solution at 37%). After fixation, samples were decalcified for 2 days with a solution of 2% ascorbic acid and 0.3 M sodium chloride. After complete decalcification, samples were dehydrated for 24 h in graded ethanol 70° to 95° followed by a bath of butanol for 24 h at 60°C and finally embedded in paraffin (Paraplast Plus). Five µm-thick sections of arm and calyx samples were cut with a Microm HM340 E microtome and mounted on silane-coated glass slides.

#### Staining and imaging

Transverse sections of the calyx and longitudinal sections of the pinnules were stained with Masson’s trichrome stain. Images were taken using a microscope (Orthoplan optical microscope, Leica) equipped with a high sensibility camera (DFT7000 T, Leica) using the Application Suite X software (LAS X, Leica).

### Scanning electron microscopy

Two crinoid specimens were prepared (calyx and arms separated) for precise anatomical observation under a scanning electron microscope. After fixation with non-acetic Bouin’s fluid, the specimens were first progressively dehydrated with ethanol before being dried in a chamber using the CO_2_ critical point drying technique. The samples were then coated with a thin film of gold and palladium using a Jeol JFC-1100E sputter coater (JEOL Company, Tokyo, Japan). Observations of the fine structure of the pinnules and calyx of the feather star were carried out with a Jeol JSM-7200F scanning electron microscope.

### Immunodetection of opsins on crinoid tissue sections

To detect opsins within the tissues of *A. bifida*, both immunofluorescence and immunohistochemical methods were carried out using two sets of purified polyclonal antibodies. These two primary antibodies, anti-Arub_ops1.1 and anti-Arub_ops4, produced in rabbits by Eurogentec, are directed respectively against ciliary opsin 1.1 (residues 159 to 174) and the c-terminal part of rhabdomeric opsin 4 (residues 447 to 461) (**Suppl. Fig. S2**) from the common European sea star species *Asterias rubens*. Histological paraffin sections of the calyx and pinnules of *A. bifida* specimens were used for opsin immunodetection.

#### Immunofluorescence

The dewaxed and rehydrated sample sections were rinsed with a solution of phosphate-buffered saline containing 0.5% Tween (PBS-T) and blocked 1h in PBS-T with 3% BSA (Bovine Serum Albumin). Samples were then incubated overnight at 4°C with either primary anti-opsin 1 antibodies or anti-opsin 4 antibodies at 1:200 dilution. Tissues were then rinsed in PBS-T and incubated 1h with commercial secondary antibodies (goat anti-rabbit IgG coupled to a Texas-Red fluorochrome; Alexa-fluorTM 594, Invitrogen A11012) at 1:100 dilution, revealing red fluorescence (peak at 618 nm) after exposure to an optimum excitation length of 590nm. After a final rinse with PBS-T, the slides were mounted with an aqueous mounting medium (Vectashield, Vector Laboratories) containing DAPI (4’,6-diamidino-2-phenylindole) highlighting cell nuclei with blue fluorescence (450-490 nm). Opsin labelling was visualised by an Olympus FV1000D inverted confocal laser scanning microscope under three channels of fluorescence by laser excitation (peak excitation/emission): DAPI (358/461 nm), FITC (495/521 nm) and Texas Red (590/618 nm).

#### Immunohistochemistry

Sections used for immunohistochemistry were treated slightly differently from those used for immunofluorescence. After dewaxing and rehydration, slides were heated in a citrate buffer (0.01 M, pH 6.2) in the microwave (2X5 min, 900 W) to unmask specific antigenic epitopes present in the tissue. Tissues were then immersed in 0.06% hydrogen peroxide to quench endogenous peroxidases. The following steps of rinsing, blocking and incubation treatments with primary antibodies were similar to those described above for immunofluorescence. After a second rinse, the tissues were incubated 1 h with a horse anti-rabbit/peroxidase secondary antibody complex (ImmPressTMReagent kit, Vector Laboratories). For opsin revelation, sections were incubated 1 to 3 minutes with a PBS-buffered solution (pH 7.2) containing 4 mg/ml DAB (4-dimethylaminoazobenzene) and 0.02% H_2_O_2_. The sections were counterstained with hemalun/Luxol Blue to enhance the contrast with the brown DAB-immunostaining.

#### Controls

Various negative and positive controls were conducted. Negative controls included omitting the primary antibody or using a pre-immune rabbit serum diluted at 1:2000 in place of the primary antibody. In each case, all control sections were not labelled **(Suppl. Fig. S3)**. Sections from tissues of *Asterias rubens*, including the optic cushions, tube feet and nervous system were used as positive controls for the detection of opsins 1 and 4. In all cases, immunostaining yielded the expected results, as described in previous publications [Ullrich-Lüter et al. 2011, Clarke et al. 2024].

## Results

### *Antedon bifida* is negatively phototactic with a maximum response under blue light

*Behavioural tests revealed that adult A. bifida crinoids exhibited a significant negative phototactic response to white light (p = 0.0099*\*\*), moving away from the source compared to controls without light. **(Fig. 2B)**. This negative phototactic behaviour was also significant with the red light (p = 0.031*) and reached its maximum amplitude with blue light (p = 0.0062**). This result points to a high sensitivity of crinoids to the blue part of the visible spectrum. In contrast, the results for the green-light tests showed no significant difference in displacement values compared to the dark control values (green: p = 0.167).

Displacement values were normalised by calyx size to account for individual size variation. However, we found that there was no correlation between the size of the individuals and the travelled distance during the test. After normalisation, significant differences were only observed for the blue and white lights compared to the dark control.

Absolute displacement values were also considered (non-oriented displacements) to determine the reactivity of crinoid individuals to light without considering the direction taken **(Fig. 2C)**. As illustrated in Figure 2C, the distances travelled during the test, independently of the direction, did not vary significantly between individuals exposed to the different wavelengths and the controls remaining in the dark.

Thus, although light stimuli did not induce significant changes in the mobility of *A. bifida* (non-directional displacement), white, red (630 nm) and blue (463 nm) light stimuli induced a significant directional displacement of crinoids away from the light source (negative phototactic behaviour).

### *Antedon bifida* has a reduced opsin repertoire composed of only three rhabdomeric opsins

The homology-based opsin search in the *A. bifida* genome enabled the discovery of three opsin genes: the *Abif-opsin 4.1* gene present on chromosome 4 [NCBI reference: OY727096.1], the *Abif-opsin 4.2* and *Abif-opsin 4.3* genes present on chromosome 6 [NCBI reference: OY727098.1] **(Fig. 3B)**. The three predicted *A. bifida* opsins contain, respectively 417, 356 and 408 amino acids (46.42, 41.17 and 45.63 kDa). A clue as to the identification of these three sequences was obtained through reciprocal blast searches and comparison of the sequence similarity with the set of reference opsins present in the sea urchin *S. purpuratus* and three other non-echinoderm opsin types **(Fig. 3A)**. The results showed that the three *A. bifida* opsin sequences have higher similarity with the (non-canonical) rhabdomeric opsins of echinoderms (opsin 4). The same analyses carried out on the genome of another closely related crinoid species, *Nesometra sesokonis*, also revealed the presence of three similar opsin sequences, which also appeared to belong to the rhabdomeric opsin group. On the other hand, research carried out on the genome of the species *Anneissia japonica* (Comatulidae) revealed no opsin genes. It therefore seems that only one bilateral opsin lineage is still present in *A. bifida* and presumably in all Crinoids of the family Antedonidae.

**Figure 3.**
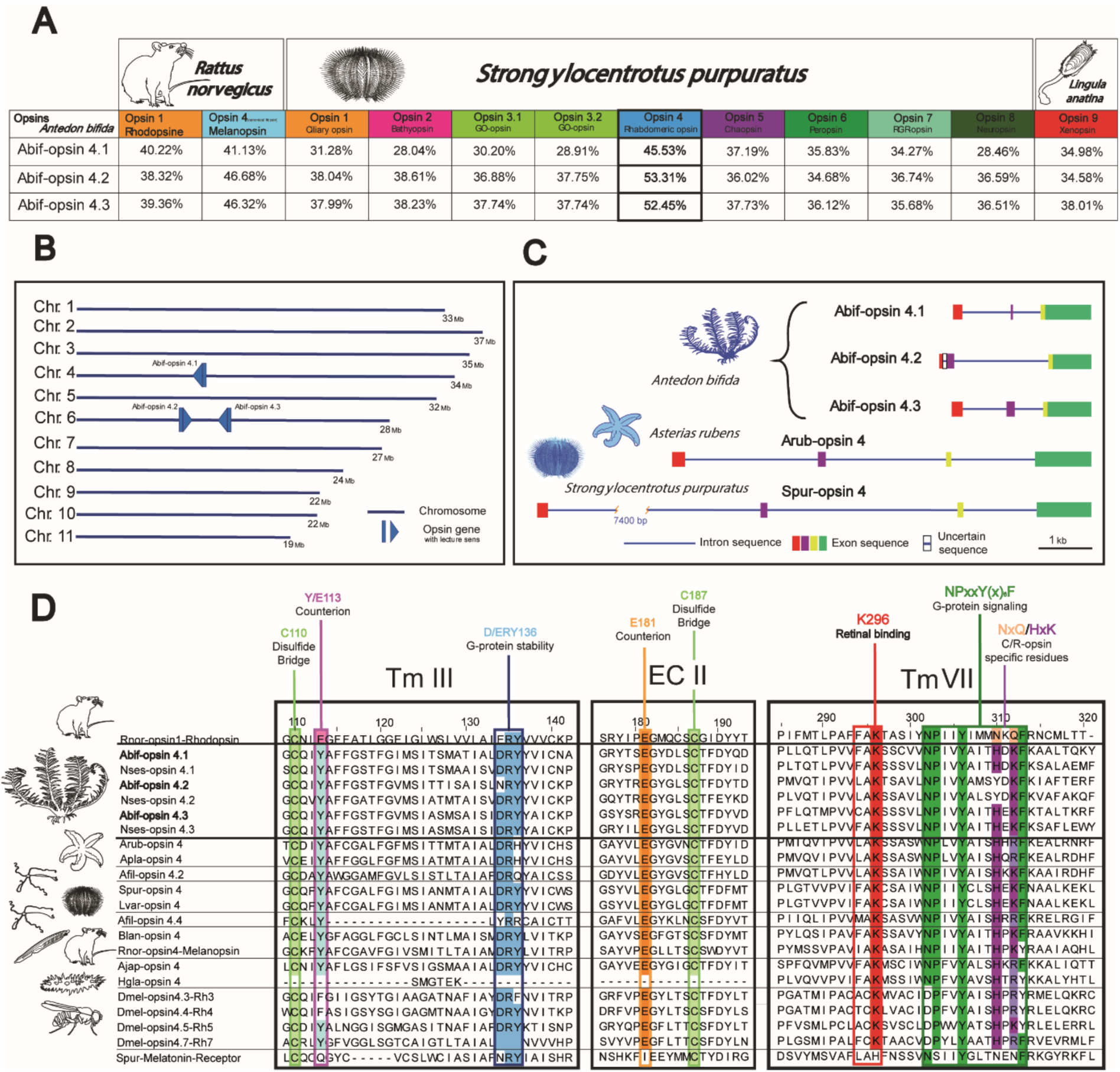
*In silico* analysis of crinoid opsins: (A) Similarity comparison (%) between the three crinoid opsin protein sequences predicted from the *Antedon bifida* genome with all echinoderm opsin groups present in the model species of sea urchin (*Strongylocentrotus purpuratus*). This comparative study was also extended to two vertebrate canonic opsins present in *Rattus norvegicus* and one non-echinoderm opsin group in a brachiopod *Lingula anatina*. The three bold boxes highlight the best similarity percentage between the crinoid opsins and the non-canonical rhabdomeric opsin of *S. purpuratus*. (B) The chromosomal position of the three opsin genes in the *Antedon bifida* genome represented by a leading arrow indicating the reading direction of the gene. (C) The representation of intronic and exonic regions (and indication of homologous regions) in the three opsin genes of the studied species (*A. bifida*) compared to the sea star and sea urchin rhabdomeric opsin genes of the model species (*A. rubens* and *S. purpuratus*). (D) Alignments in the third (Tm III) and seventh (Tm VII) transmembrane helix domains as well as in the second extracellular loop (EC II) protein region of the six opsin sequences evidenced in both crinoid species (Abif = *Antedon bifida*; Nses = *Nesometra sesokonis*) versus 14 rhabdomeric opsins of other bilaterian metazoans (Arub = *Asterias rubens*; Apla = *Acanthaster plancii*; Spur = *Strongylocentrotus purpuratus*; Lvar = *Lytechinus variegatus*; Afil = *Amphiura filiformis*; Blan = *Branchiostoma lanceolatum*; Rnor = *Rattus norvegicus*; Ajap = *Apostichopus japonicus*; Hgla = *Holothuria glaberrima*; Dmel = *Drosophila melanogaster*). The first and the last supplementary sequences in these alignments are respectively the rat rhodopsin (a reference ciliary opsin which gives the amino acid numeration) and the melatonin receptor of sea urchin *S. purpuratus* (non-opsin GPCR protein) used as comparison models. Different amino acid patterns, characteristic of typical opsins, are highlighted in colour in these alignments and are all present in the three crinoid opsins.

All *Abif-opsin* genes share a conserved gene structure composed of one or two short exon sequences (80-270 bp) followed by a third, much longer, terminal exon region (800-960 bp) **(Fig. 3C)**. In addition, our genomic analysis of *A. bifida* opsin genes revealed two major structural differences compared to rhabdomeric opsin genes in other echinoderm classes, which lead to an overall shorter gene length due to a lower number of intronic regions (between 1 or 2 introns for *A. bifida* versus 3 for sea stars and sea urchins) and reduced intron sizes (between 540 and 1,800 bp maximum for crinoid opsins and from minimum 2,300 bp for the sea star *A. rubens* to maximum 11,400 bp for the sea urchin *S. purpuratus*). Genomic sequence alignments showed homology at the level of the first and last long exon for the three crinoid opsins, as well as for those of sea star and sea urchins. Finally, in the sea star *A. rubens* and the sea urchin *S. purpuratus* opsin 4 orthologs, the regions homologous to the third and fourth exon are fused into a single last exon in the three crinoid opsin sequences. Sequence homology analysis also shows that the first exon of *A. bifida* opsin 4.2 is composed of the assembly of the two first exons homologous to those present in the other two crinoid opsins. A short region of 57 base pairs is also found between these exons, and whose exonic nature recognised by gene prediction software is still uncertain.

Multiple alignment of the predicted opsins found in *A. bifida* and *Nesometra sesokonis* with other rhabdomeric opsins reveals the presence of highly conserved amino acid residues characteristic of light-sensitive opsins **(Fig. 3D)**. These opsin characteristics are mainly amino-acids present in the third and seventh transmembrane helix (Tm) as well as in the second extracellular loop (EC), which allow interactions with the conjugated chromophore molecule (retinal) and the G-protein. The most representative opsin residue is a lysine residue in the 7^th^ Tm, corresponding to K296 position in the rat rhodopsin, which performs the Schiff-based covalent binding with the retinal [Terakita 2005]. This lysine is conserved in all aligned predicted opsins of crinoids. For the stabilisation of this protonated Schiff-based binding, a counterion is necessary and this function is typically assumed by a glutamate residue [Terakita et al. 2004; Varma et al. 2019]. The probable ancestral position of this counterion is the equivalent of the glutamate E181 in the rat rhodopsin present in the second extracellular loop. This E_181_, as the only counterion function, is typical of bistable opsins such as rhabdomeric opsins [Terakita et al. 2012; 2014]. The monostable opsins like vertebrate ciliary opsins possess another glutamate counterion in Tm III (corresponding to the E113 in the rat rhodopsin) closer to the retinal binding. This glutamate E_113_ is replaced by a tyrosine residue in rhabdomeric opsins. The three crinoid opsins present the same ancestral counterion glutamate E181, with the tyrosine residue also replacing E113 like in other echinoderm rhabdomeric opsins. These results point to the probable bistable character of crinoid opsins. Another typical amino acid pattern of all GPCR proteins like opsins is the NPxxY(x)_6_F pattern (corresponding to the sequence NPII(Y)306 in the rat rhodopsin) which could contribute, amongst other functions, to the binding between the opsin and the G-protein [Fritze et al. 2003]. This motif is also present in all the sequences including the Melatonin GPCR receptor. Another tripeptide also seems to participate in the association between the opsin and the G-protein, this is the (D)RY motif (corresponding to the ER(Y)136 in the rat rhodopsin). We can also mention the two opsin-typical cysteine residues (corresponding to C110 in the Tm III and the C187 in the EC II of rat rhodopsin) which allow the stabilisation of protein conformation owing to the formation of a disulfide bridge. The three crinoid opsins possess all these key residues typical of functional opsins. We can also note the presence, in two of the three crinoid opsins (4.1 and 4.3), of the HxK pattern inside the motif NPxxY(x)6F, which is rather typical of rhabdomeric opsins and is homologous of the pattern NxQ found in ciliary opsins (corresponding to the sequence Nx(Q)312 in the rat rhodopsin) [Upton et al. 2021]. This typical-rhabdomeric pattern is only partly present in the crinoid-opsins 4.2 in which the histidine residue is replaced by a tyrosine. It is interesting to note that this is the only case in the echinoderm rhabdomeric opsins identified thus far.

### Phylogenetic position of crinoid opsins

A first exploded global opsin phylogeny in different species of Echinoderms and other typical bilaterian phyla allowed it to recover all nine ancestral bilaterian opsin lineages **(Fig. 4A, full phylogenetic tree presented in Supplementary Figure S1)**. In this way, we were able to confirm in *silico* analyses that all three crinoid opsins belong to the rhabdomeric opsin group (opsins 4). A focus on rhabdomeric opsins 4 then allowed us to highlight their relationships between investigated crinoid species and other echinoderms **(Fig. 4B)**. Crinoid and other echinoderm rhabdomeric opsins gathered into a well-separated clade **(Fig. 4B)** that further branched apart from classical rhabdomeric opsins like vertebrate melanopsins and *Drosophila* r-opsins, supporting that all rhabdomeric opsins in echinoderms diversified from a unique common ancestor. This is consistent with the fact that all echinoderm opsins 4 have so far been identified as non-canonical rhabdomeric opsins and in this phylogeny, appear to be the sister group to the canonical opsins of the chordates and protostomes. Other “non-conventional” rhabdomeric opsins of brachiopods and polychaetes are not directly grouped with echinoderm opsins and are represented as a sister group of the clade containing canonical opsins and non-canonical opsin of echinoderms. It is also important to notice that the three opsin sequences discovered from the genomes of both crinoid species (*Antedon bifida* and *Nesometra sesokonis*) are exclusively grouped with two other published partial opsin sequences from the transcriptomes of two other feather star species. This crinoid opsin clade is placed in the basal position of the other echinoderm rhabdomeric opsins, in agreement with the phylogeny of extant echinoderms. The other echinoderm opsins 4 are also grouped according to the different echinoderm classes, in perfect agreement with the well-established phylogenetic classification of the phylum. The relationships between crinoid opsins point to the existence of three rhabdomeric opsin lineages in the Antedonidae family, the most basal position being occupied by crinoid opsin 4.2 orthologs.

**Figure 4.**
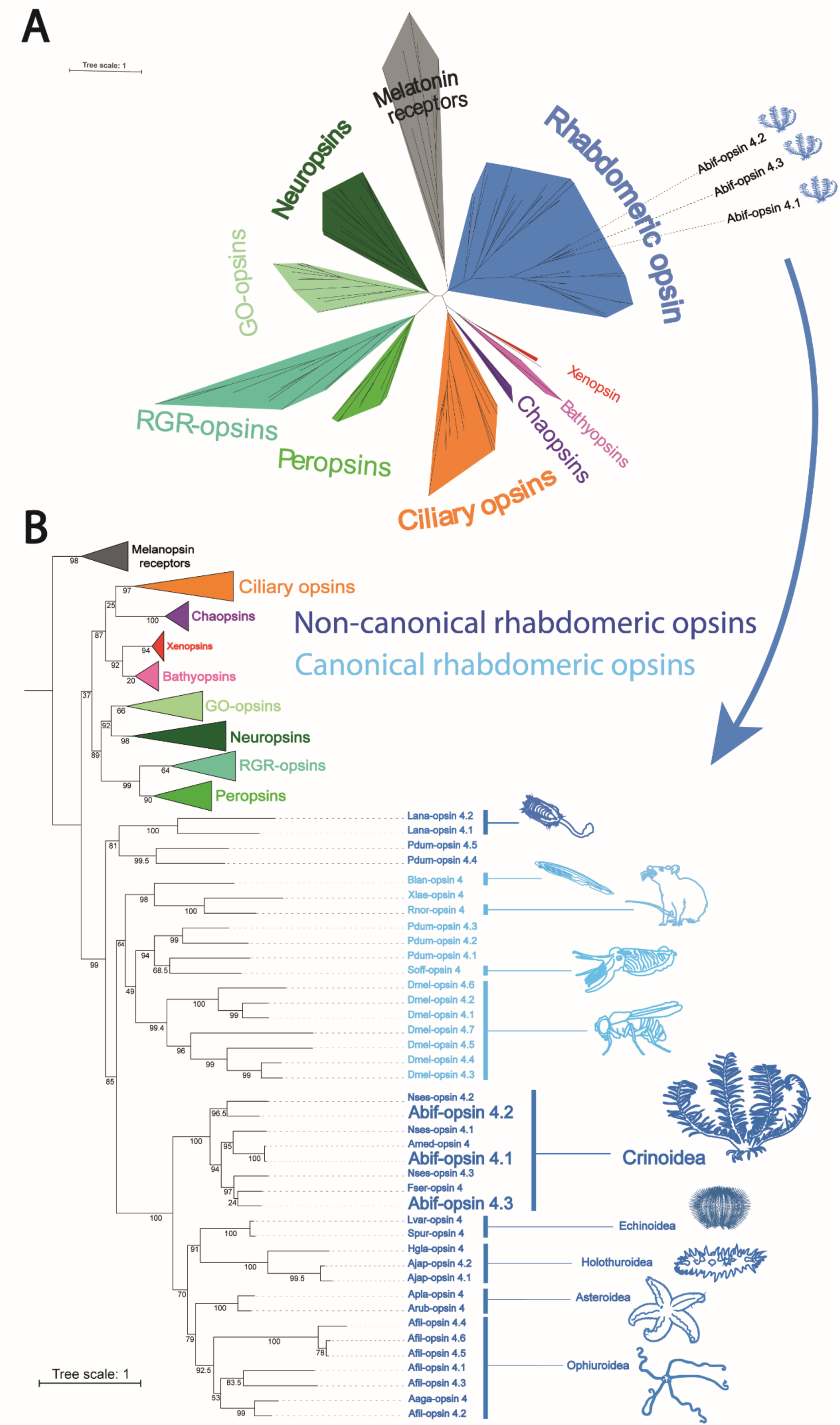
Phylogenetic analysis of Crinoid opsins. (A) Global phylogenetic tree of the nine representative ancestral opsin lineages in bilaterian metazoans supported by maximum likelihood method. Bootstrap values are indicated at the root of each opsin cluster. The melatonin receptor cluster is used to root the phylogenetic tree and the branch length scale gives an insight into the relative rate of amino acid substitution per site. The three opsin sequences evidenced in the crinoid *Antedon bifida* (Abif) are indicated in bold into the rhabdomeric opsin cluster and associated with blue crinoid icons. (B) Focus on the rhabdomeric opsin cluster with a distinction between Canonical (in light blue) and non-canonical (in dark blue) R-opsins. The branch lengths and bootstrap values are indicated in this case respectively above and below the branches. There are in total 40 rhabdomeric opsin sequences present in this phylogeny belonging to 18 representative bilaterian species (Echinoderms: Abif = *Antedon bifida*; Amed = *Antedon mediterranea*; Nses = *Nesometra sesokonis*; Fser = *Florometra serratissima*; Arub = *Asterias rubens*; Apla = *Acanthaster plancii*; Spur = *Strongylocentrotus purpuratus*; Lvar = *Lytechinus variegatus*; Afil = *Amphiura filiformis*; Aaga *= Astrotomma agassizii*; Ajap = *Apostichopus japonicus*; Hgla = *Holothuria glaberima*; Chordates: Blan = *Branchiostoma lanceolatum*; Rnor = *Rattus norvegicus*; Xlae = *Xenopus laevis*; Lophotrochozoans: Lana = *Lingula anatina*; Pdum = *Platynereis dumerilii*; Soff = *Sepia officinalis*; Arthropods: Dmel = *Drosophila melanogaster*). The three opsin sequences of the crinoid species (*A. bifida*) (Abif-opsin 4.1, 4.2 and 4.3) are highlighted in bold at the base of the non-canonical rhabdomeric opsin cluster of Echinoderms. Accession numbers of all sequences are presented in the Supplementary Table S1. The full opsin phylogeny is presented in the Supplementary Figure S1.

### Crinoid opsins are functional photosensitive receptors

#### Structural modelling of protein photoreceptors

Homology modelling indicates overall close structural relatedness between all three opsins of *Antedon bifida* (Abif-opsin) despite a shorter transmembrane helix 1 (H1) for Abif-opsin 4.2 **(Fig. 5B)**.

**Figure 5.**
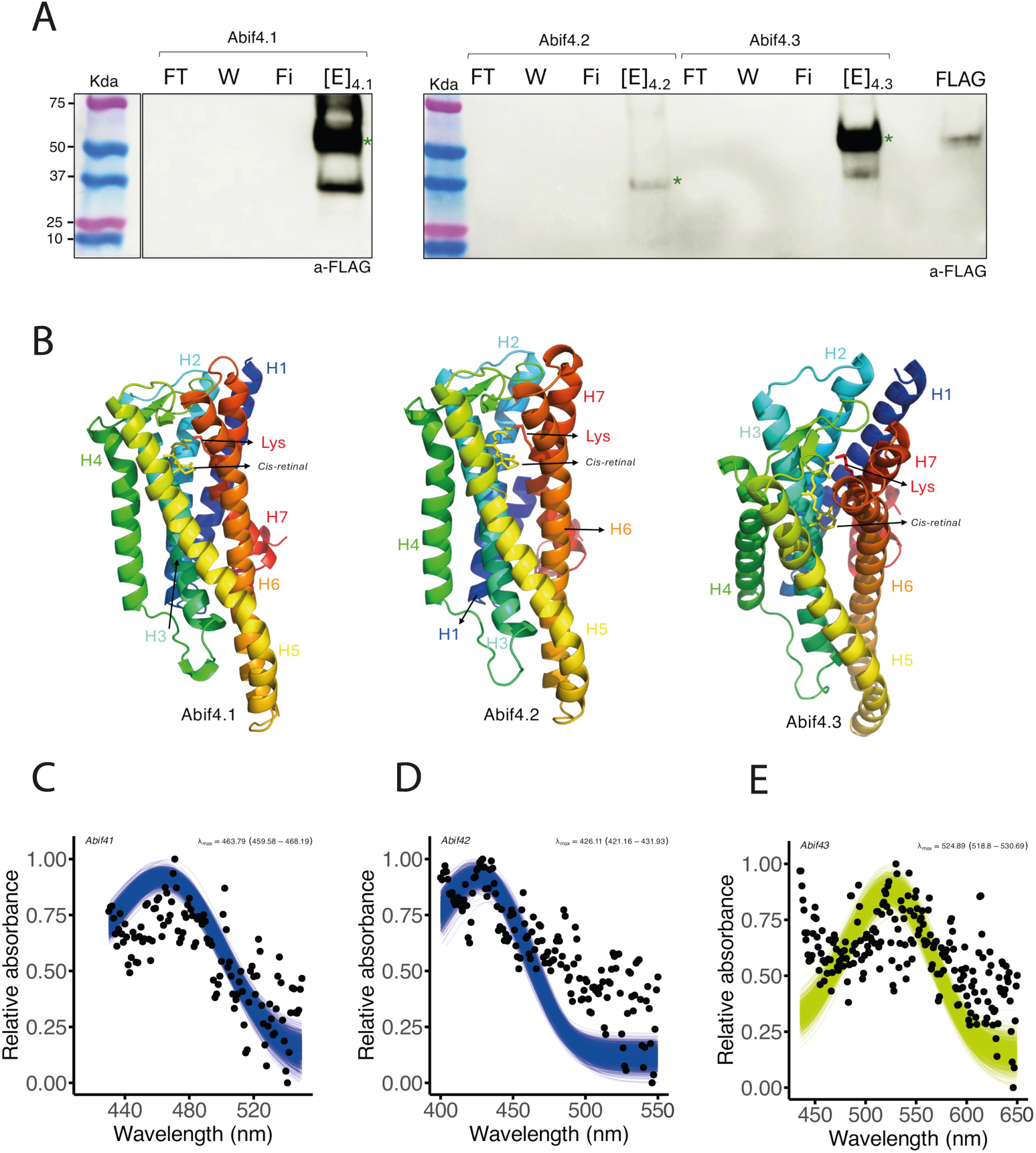
*In vitro* characterisation of photosensory function and light absorption of the three crinoid opsins (Abif-opsins). (A) Immunoblot analysis of purified photoreceptor complexes (Abif-opsins, 11-cis retinal and a FLAG recognising peptide) present after their heterologous expression in HEK293T. Gel migration shows a band for the concentrate eluate ([E]) at the relative expected molecular weight for each of the three opsins expressed *in vitro* (Abif 4.1: 49.73 kDa; Abif 4.2: 40.99 kDa and Abif 4.3: 49.19 kDa) indicated by a green asterisk. The last loaded well (FLAG) correspond to a positive control test using FLAG proteins (50 kDa). (B) Modelling and comparison of the three-dimensional structure of the three Abif-opsins. Except the shorter helix 1 (H1) of the Abif 4.2, the strong structural resemblance of the three crinoid opsins confirms that they are closely related to each other. (C-E) Light absorbance spectra recording by UV-vis spectroscopy (dark dots= mean relative absorbance) and analysis by an *in silico* polynomial visual template, for each Abif-opsins expressed *in vitro* in presence of 11 cis-retinal. (C) Absorbance spectrum of Abif opsin 4.1 (459-468 nm) corresponding to the blue light with λmax = 463 nm. (D) Absorbance spectrum of Abif opsin 4.2 (410-500 nm) corresponding also to the blue light with λmax = 426 nm. (E) Absorbance spectrum of Abif opsin 4.3 (450-600 nm) corresponding to the green light with λmax = 524 nm. FT= flow through; W= wash; E= eluate; Fi= filtrate; [E]= concentrate eluate, FLAG= positive control.

#### In vitro light absorption of crinoid opsins

The purified *A. bifida* opsins expressed heterologously in HEK293T cells formed active complexes in solution in the presence of 11, *cis*-retinal. Although Abif-opsin 4.2 appeared more moderately expressed, the eluted Abif-opsin 4.1 and Abif-opsin 4.3 protein fractions were highly enriched **(Fig. 5A)** consistent with proper cellular expression and trafficking to the plasma membrane. Abif-opsin 4.1 absorbed wavelengths of light ranging from 440 nm to 520 nm with a maximal relative absorption around 464 nm (lambda max) with confidence intervals from 459 to 468 nm **(Fig. 5C)**. Abif-opsin 4.2 absorbed wavelengths of light ranging from 410 nm to 500 nm with a maximal relative absorption around 426 nm with confidence intervals from 421 to 432 nm **(Fig. 5D)**. Abif-opsin 4.3 absorbed wavelengths of light ranging from 450 nm to 600 nm with a maximal relative absorption around 524 nm with confidence intervals from 518 to 530 nm **(Fig. 5E)**. These results support that all three encoded opsins are functional to absorb short and medium wavelengths.

### *Antedon bifida* opsins are expressed in the tube feet and nervous systems

#### Opsin detection in the calyx area

*A. bifida* has a typical morphology of feather star crinoids (*Comatulida*) with a central calyx with mobile hook-like cirri on the aboral side for locomotion **(Fig. 6A)** and ten filtering arms connected to the oral side of the calyx. Each arm possesses numerous bilateral ramifications called pinnules **(Fig. 6B)**. A deep aboral nervous system, typical of crinoids, is localised in the deep part of the calyx and constituted by a complex bowl-shaped cluster whose branches form the brachial nerves **(Fig. 6F)**. Ramifications of brachial nerves extend into the aboral side of the pinnules. The mouth is in the middle of the calyx oral side (tegmen), right next to the anal tube that is slightly off-centre **(Fig. 6C)**. Five ambulacral grooves, that carry filtered food to the mouth, start from the mouth and dichotomise before extending to the arms. Each calyx groove is alternately bordered on both sides by spherical bodies called saccules (**Fig. 6D-F)** and by a unique series of tube feet **(Fig. 6E-F)**. Histological cross-sections of the ambulacral groove region point to the presence of two neuronal networks, on one hand, the ectoneural basiepithelial nerve plexus confined at the base of the groove epidermis and, on the other hand, a diffuse hyponeural nerve plexus present slightly down in the connective tissue **(Fig. 6F)**.

Both ecto- and hyponeuronal plexus exhibited a strong immunoreactivity after exposure to antibodies raised against opsin 4 of the sea star *Asterias rubens*, supporting the presence of opsins in these nervous structures **(Fig. 7 A-E)**. The high sequence similarity between the antigen sequence of *A. rubens* opsin 4 and the sequence of *A. bifida* opsin 4.1 **(*Suppl. Fig. S2*)** suggests that the immunoreactivity observed both by immunofluorescence **(Fig. 7 A-C)** and by immunohistochemistry **(Fig. 7 B-E)** correspond to the expression of the Abif-opsin 4.1 in these nervous structures.

**Figure 6.**
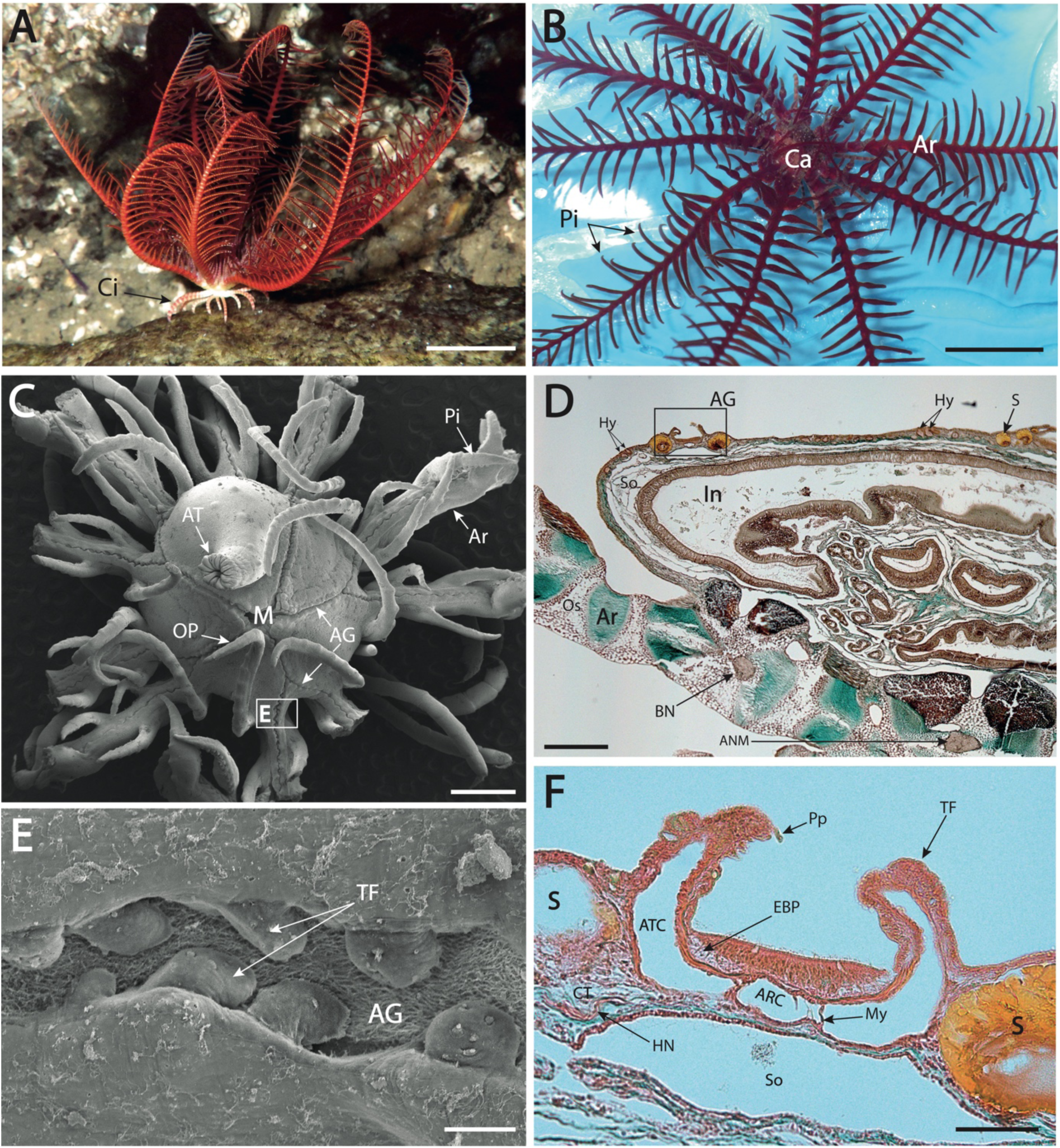
Morphological structure of the Calyx: (A) An adult specimen of *Antedon bifida*. (B) Oral view of the central calyx surrounded by the ten branched arms. (C) Scanning electron microscopy picture of the whole oral surface (tegmen) of the calyx with a focus on the ambulacral groove indicated by a box. (D) Lateral histological cross section on the global calyx with a focus on the boxed ambulacral groove region. (E) Micrography of the calycinal groove portion bordered by well-visible tube feet. (F) Transversal histological section (Masson’s trichrome stain) of the ambulacral groove on the calyx. AG: Ambulacral Groove; ANM: Aboral Nerve Mass; Ar: Arm; ARC : Ambulacral Radial Canal; AT: Anal Tube; ATC: Ambulacral Tentacular Canal; BN: Brachial Nerve; Ca: Calyx; Ci: Cirri; CT: Connective Tissue; EBP: Ectoneural Basiepithelial Plexus; HN : Hyponeural Nerve; Hy: Hydropores; In: Intestine; M: Mouth; My: Myocyte; OP: Oral Pinnule; Os: Ossicule; Pi: Pinnules; Pp: Papilla; S: Saccule; So: Somatocoele; TF: Tube Foot. Scales: A. 20 mm, B. 10 mm, C. 400 µm, D. 300 µm, E. 20 µm, F. 50 µm.

**Figure 7.**
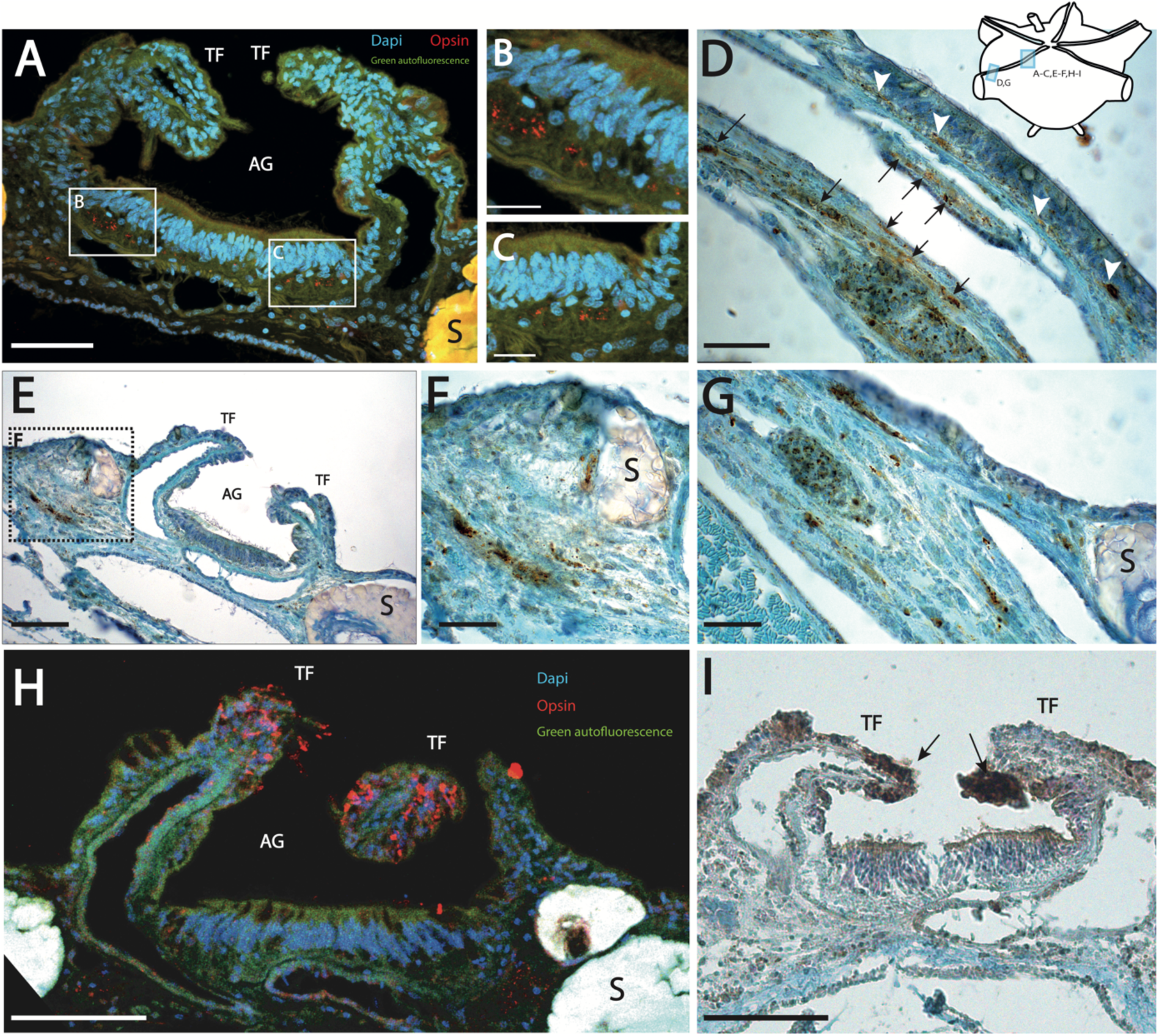
Immunostainings of two rhabdomeric opsins in the Calix ambulacral groove tissues of *Antedon bifida*: (A) Localisation of opsin 4.1 by red immunofluorescence into the ectoneural basiepithelial nerve plexus just under the ambulacral groove epithelium. (AI; AII) Focus on the two basiepithelial spots of opsin labelling in red. (B) Opsin 4.1 is highlighted at the base of arm with a localisation on the one hand in the ectoneural basiepithelial plexus (white head arrow) and the other hand in the hyponeural plexus present deeper into the connective tissue (black arrow). (C) Localisation of the same opsin 4.1 by immunocytochemistry in brown into the hyponeural nerve plexus inside the connective tissue of the calyx ambulacral groove. (D) Focus on the hyponeural opsin showing the localisation near a saccule. (E) Focus on the brown immunostaining of opsin 4.1 in the hyponeural plexus of the base of the arm. (F) Labelling in red by immunofluorescence of opsin 4.2 at the top of tube feet (in the sensory papillae) bordered the ambulacral groove of the calyx. (G) Immunocytochemistry showing the opsin 4.2 expression at the top of tube feet (arrow). Scales: A. 50µm, A’-A’’. 10µm, B. 25µm, C. 50µm, D. 20µm, E. 20µm, F. 50µm, G. 50µm.

Other structures showing the presence of opsins corresponded to tube feet, bordering the ambulacral grooves **(Fig. 7 E-F)**. The epidermis of each tube foot presents several papillae with ciliary sensory cells. The opsin immunoreactivity was observed more precisely in the epidermal cells at the base and tip of these sensory papillae **(Fig. 7 F-G)**. These immunodetections were performed with antibodies raised against *A. rubens* opsin 1.1. The similarity of antigenic sequence between *A. rubens* opsin 1.1. and *A. bifida* opsin 4.2 **(Suppl. Fig. S2)** suggests that the immunoreactivity observed in the papillae of tube feet corresponds to the expression of the Abif-opsin 4.2. However, sequence divergence for the target binding region in the available peptide antibodies does not rule out that opsin 4.3 may co-localise with opsin 4.2 or be expressed in additional sensory cells.

#### Opsin detection in the pinnules

The pinnules of crinoids are lateral extensions of the arms **(Fig. 8A)**. A pinnular ambulacral groove occupies the middle oral side of each pinnule and is bordered by two rows of tube feet allowing food capture and transport **(Fig. 8B)**. The pinnular tube feet are grouped in triads arranged alternatively on either side of the groove **(Fig. 8C)**. These triads consist of one long primary tube foot and a medium-sized secondary tube foot dedicated to trap food particles, and a short tertiary tube foot whose function is to form the food bolus that is transported to the mouth [Lahaye and Jangoux 1985]. Similarly to the calyx tube feet, pinnular tube feet also have several epidermal sensory-secretory outgrowths called papillae. Papillae are known to feature several cell types: secretory cells, peripheral support cells with a long apical cilium, and central neurosensory cells with a shorter cilium [Flammang and Jangoux 1991]. Scanning electron microscopy observations highlighted ciliary structures at the extremity of the papillae **(Fig. 8H)**. As for the case in calyx papillae, these pinnular papillae showed an immunoreactivity with the anti-Arub_opsin 1.1 antibodies **(Fig. 8 E-G)**. These opsin labelling corresponds more probably to the expression of Abif-opsin 4.2 as mentioned above (***Suppl. Fig. S2***). The labelling was strong in the basal part of papillae but was also present occasionally in the apical pole of papillae cells **(Fig. 8 E-G)**.

**Figure 8.**
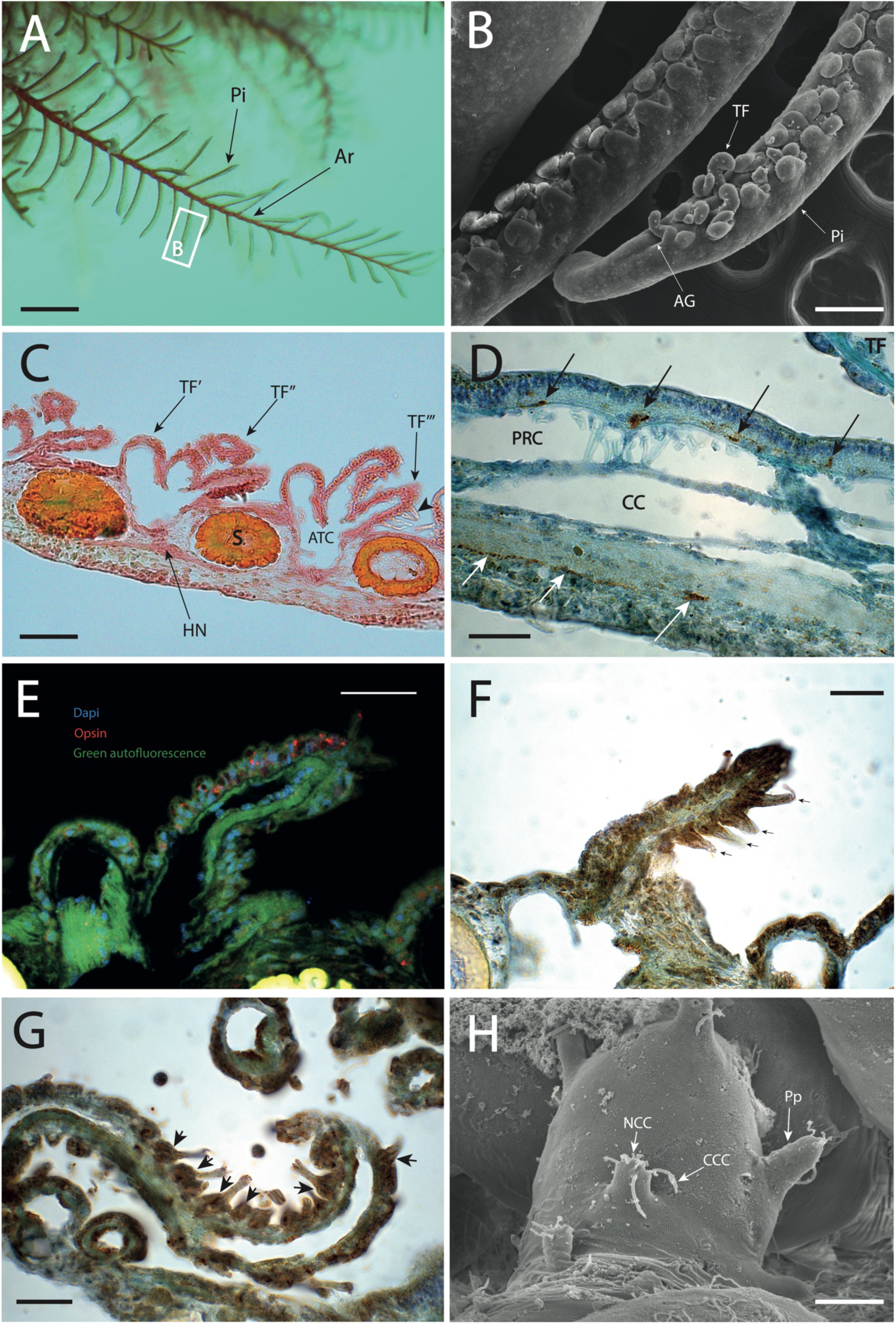
Anatomical structures of pinnules tissues and rhabdomeric opsin immunostainings in the pineal ambulacral groove in *Antedon bifida*: (A) Crinoid arm with a focus on a pinnule indicated by a red box. (B) Scanning electron microscopy picture at the extremity of the pinnule. (C) Longitudinal histological section (Masson’s trichrome stain) in the lateral pinnule edge. (D) Localisation of opsin 4.1 in the ectoneural basiepithelial plexus (dark arrow) and the deep aboral nerve of the pinnule (white arrow). These histological structures become visible thanks to the further longitudinal section in the centre of the ambulacral groove. (E) Labelling of opsin 4.2 by immunofluorescence in red into the tube feet epidermis and the extremity of sensory papillae. (F) Immunostaining (brown precipitate) of opsin 4.2 in the lateral sensory papillae (dark arrow) of one secondary tube foot. (G) Similar opsin 4.2 immunostainings in brown at the base of sensory papillae (dark arrow) of one primary tube foot. (H) Micrography of the sensory papillae structures of a tube foot. AG: Ambulacral groove; Ar: Arm; ATC: Ambulacral Tentacular Canal; CC: Coelomic Canal; CCC: Covering ciliary cell; NCC: Neuronal Ciliary Cell; Pi: Pinnules; TF: Tube foot; TF’/TF’’/TF’’’: primary/secondary/tertiary Tube Foot; HN: Hyponeural nerve; Pp: Papilla; PRC: Pinnule Radial Canal. Scales: A. 5mm, B. 200µm, C. 50µm, D. 20µm, E. 20µm, F. 20µm, G. 20µm, H. 3µm.

The three parts of the pinnular nervous system are extensions of those found in the arms and calyx. The pinnular ambulacral grooves feature an ectoneural basiepithelial plexus and a deep aboral nerve plexus in the centre of the pinnules. On the other hand, the hyponeural nerve plexus is constituted by two lateral nerve cords that run longitudinally on either side of the pinnules and innervate the tube feet and saccules **(Fig. 8 C)**. As was the case in the calyx section, aboral, ectoneural and hyponeural networks present in pinnules expressed opsin proteins **(Fig. 8D)**. As mentioned before this immunostaining in nervous structures corresponds, more probably, to the presence of Abif-opsin 4.1 (**Suppl. Fig. S2**).

## Discussion

### Crinoid light behaviour

Many reef-dwelling Comatulid species exhibit photophobic behaviours, feeding primarily at night and hiding in rocky crevices or folding their arms and pinnules during the day [Magnus 1962; Meyer 1973; Rutman and Fishelson 1969]. Our results evidenced a negative phototactic behaviour in *Antedon bifida* which is consistent with previous *in situ* observations of hiding behaviour made in other shallow water feather stars. One study also mentioned a positive phototactic behaviour for a small feather star species *Dorometra nana* that was attracted to a light source [Clark 1909]. This isolated observation could indicate a species-specific response, but the photic response could also be affected by different variables such as the light intensity. Some observations suggested that the nocturnal feeding activity and the diurnal shelter-seeking behaviour of many crinoids are not linked to the circadian cycle but would be well induced by direct ambient light detection. Indeed, some specimens belonging to the same species (*Heterometra savignii)* but living at greater depths, in an environment of constant low luminosity, tended rather to lose their nocturnal activity but kept their high photophobic reaction to an intense light stimulus [Fishelson 1974]. It is therefore not surprising that the studied population of *A. bifida* living just below the surface, clinging to the shade under pontoons of the marina, flees the polychromatic white light as it was previously reported in the literature based on observations in natural conditions [Dimelow 1958].

Behavioural tests on *Antedon bifida* revealed a pronounced negative phototactic response to blue light, indicating higher sensitivity to shorter wavelengths (λmax = 463 nm). The greater sensitivity to blue light has also been demonstrated several times by phototactic behavioural tests in other echinoderm groups, such as the sea urchin *Strongylocentrotus purpuratus* [Ullrich-Lüter et al 2011], the brittle star *Amphiura filiformis* [Delroisse et al 2014] and, more recently, the sea cucumber *Holothuria leucospilota* [Wang et al 2023]. The blue part of the light spectrum is the wavelength range that penetrates the farthest into the water column and is, therefore, the light component, together with green light, mostly present in open water environments (Davis 1991). Electrophysiological studies in ocellary structures of some reef-dwelling sea star species attested a very narrow sensitivity peak for the blue wavelength (450 nm) suggesting the presence of a blue-sensitive opsin in ocellary spots of these sea stars [Garm and Nilsson 2014, Petie et al. 2016]. By contrast, *A. bifida* is sensitive to a much broader light spectrum. Experiments performed in the present study demonstrated a significant negative phototactic behaviour for red light (λmax= 630nm). This sensitivity to red light may be correlated with the fact that this species lives at very shallow depths (less than 10m) and therefore is exposed to longer wavelengths like red penetrating only a few metres below the sea surface, especially in the Atlantic and North Sea where the light intensity declines quickly with depth (Davis 1991, Capuzzo et al. 2013, 2015). This large variability in the range of perceived light wavelengths, together with specific expression patterns, suggests that directional photoreception in this species of Comatulida may be mediated by different opsins. Our study confirms the presence of three different opsin genes evidenced in the genome of *A. bifida*, bearing all conserved characteristics of functional opsins. This suggests that *A. bifida* relies on rhabdomeric opsins for extraocular light perception. Another clue pointing to the presence of a multi-opsin photoreception in *A. bifida* is the expression of two different rhabdomeric opsins localised in specific tissues such as the subepidermal ectoneural nerve plexus and the tips of tube feet. However, further *in vitro* investigations of the bioactivity of the three different opsins are required to functionally characterise the crinoid opsins.

A trend for *A. bifida* to move away from the light source was also observed for green light (λmax = 512 nm), but this was not significant. This lack of significance is probably due either to a low detection of green light or to a lack of phototactic behavioural response for this wavelength in this Comatulid species. Green light is the wavelength range most present in the shallow coastal waters where *A. bifida* populations are found. This ecologically relevant green wavelength could therefore lead to distinct behavioural responses in contrast to other light wavelengths. A similar absence of reactivity to green light has already been described in tadpoles of the amphibian species *Xenopus laevis* living in green eutrophic water [Roberts et al. 2000]. Nevertheless, future behavioural tests with green light wavelength could be carried out on a larger number of individuals to ensure that there has been no statistical bias due to insufficient sampling.

### Opsin evolution in feather stars

*In-silico* analyses performed on the genome of the crinoid species *A. bifida* and *N. sesokonis* have revealed the presence of only three opsin genes. In the chromosome-scale genome of *A. bifida*, the opsin genes are distributed on chromosomes 4 and 6. All three opsin genes were identified as belonging to the same group of non-canonical echinoderm rhabdomeric opsins. This low opsin gene diversity was quite unexpected given that the other four echinoderm classes possess a much more varied opsin repertoire with between 9 and 15 different opsin genes currently known [D’Aniello et al. 2015]. For instance, sea urchins and brittle stars, still exhibit seven of the nine ancestral opsin lineages inherited from the common ancestor of bilaterians: ciliary opsins, bathyopsins, Go-opsins, non-canonical rhabdomeric opsins, chaopsins, peropsins/RGR-opsins and neuropsins [Ramirez et al. 2016].

Crinoids are considered as the basal lineage of echinoderms which generally justified their importance in comparative evolutionary studies. Crinoidea has a long evolutionary history of at least 485 million years [Rouse et al. 2013; Oji and Twitchett 2015] and all crinoids living today originated from the single branch of Articulata, the only crinoid lineage to have survived the great extinction of the late Paleozoic [Baumiller et al. 2011]. Current feather stars therefore represent a relatively derived group compared to the ancestral forms of the Paleozoic. The Crinoidea available genomes (i.e., *Anneissia japonica* [*ASM1163010v1* in NCBI, 0.5896 Gb], *Nesometra sesokonis* [*ASM2563120v1* in NCBI, 0.5971 Gb] and *Antedon bifida* [*ecAntBifi1.1* in NCBI, 0.3196 Gb] the only chromosome-scale genome) are comparable to sea stars (e.g., *Asterias rubens* (0.4176 Gb) and *Marthasterias glacialis* (0.52Gb) [Parey et al. 2024]) but appear considerably smaller compared to other echinoderm genomes (e.g., *Strongylocentrotus purpuratus* (0.9218 Gb), *Holothuria leucospilota* (1.4 Gb), and *Amphiura filiformis* (1.57 Gb) [Parey et al. 2024]). In *A. bifida*, specifically, a reduced number of chromosomes is observed (11 chromosomes) compared to other chromosome-scale echinoderm genomes (i.e., 18 for *Paracentrotus lividus* [Parey et al. 2024], 22 for *A. rubens*, 23 for *H. leucospilota* and 20 for *A. filiformis* [Parey et al. 2024]). Our study further reveals a reduced opsin repertoire accompanying these genomic characteristics. It is possible that, during their evolutionary history, these modern crinoids (many live at relatively great depths in low luminosity) may have lost many of the ancestral opsin groups still present in other echinoderms. The smaller crinoid genome could also explain the lower number of intronic regions in their opsin genes, making crinoid opsin genes shorter than those found in other echinoderm classes.

However, the presence of a single bilaterian opsin class may not be generalisable to all crinoid groups but may just be specific to the Comatulida family Antedonidae. Indeed, the two species (*A. bifida* and *N. sesokonis*), whose genomes have been investigated in the present study, belong to the family Antedonidae. Research carried out on the only other available crinoid genome belonging to the Comatulidae family species (*Anneissia japonica*) revealed no opsin gene. A more complete sampling across Crinoidea will be crucial to have a more complete representation of opsin diversity within the class.

The alignment comparisons with other metazoan rhabdomeric opsins have demonstrated that all three crinoid protein sequences possess the set of amino acid residues characteristic of functional opsins allowing interaction with both retinal and the G-protein. This suggests that these three rhabdomeric opsins potentially play a direct role in light detection in *A. bifida*. Our behavioural tests support the view that *A. bifida* individuals possess photosensitivity covering a wide part of the light spectrum, potentially related to the activity of different opsins exhibiting distinct spectral absorptions. The involvement of rhabdomeric opsins as the main photoreception molecular actor in the light detection has been mentioned several times in other Echinoderms [Ullrich-Lüter et al 2011; Delroisse et al 2014] notably for sea stars which possess a largely predominant expression of a rhabdomeric opsin in their compound ocelli (optic cushion) at the end of arms [Lowe et al. 2018].

The phylogenetic analysis of rhabdomeric opsins in bilaterians places the three crinoid opsins together at the base of the non-canonical rhabdomeric opsin clade of Echinoderms. Within the latter, the relationship of opsins belonging to the different classes follows perfectly the classical phylogenetic classification of echinoderms, as highlighted in other studies [Delroisse et al. 2014; D’Aniello et al. 2015; Lowe et al. 2018]. The fact that the three crinoid opsins are clustered together suggests that they originated from a duplication of an ancestral rhabdomeric opsin gene in the lineage. A similar but potentially more extensive gene duplication was observed in the brittle star species *Amphiura filiformis*, which possesses six different rhabdomeric opsin genes [Delroisse et al 2014].

### Heterologous expression of crinoid opsins confirmed their photosensory function at short light wavelengths

The probable photosensory function of these three rhabdomeric crinoid opsins discussed above is strongly supported by the heterologous in vitro expression of opsin-retinal complexes from *Antedon bifida* and their light absorption by UV-vis spectroscopy. These purified complex opsin-retinal have shown in addition a specific light absorption for the short light wavelengths: blue light for the two first opsins and green for the last one **(Fig. 9)**. This is consistent (at least for two of the three opsins) with the previous phototactic behaviour test results which indicate a higher sensitivity to blue light (the absorbance peak of the Abif-opsin 4.1 corresponding exactly to the light wavelength of maximum phototactic reaction: 463 nm). The lower but significant light red reaction (λmax= 630 nm) could therefore be the result of a broader light absorbance spectrum for one of the crinoid opsins that can potentially approximate red light (*e.g.* range of Abif-opsin 4.3= 450-600nm). On the other hand, the fact that one of the three opsins (Abif-opsin 4.3) has an absorbance in green light supports the hypothesis that green light is perceived by *A. bifida* but does not induce a significant phototactic displacement reaction.

**Figure 9.**
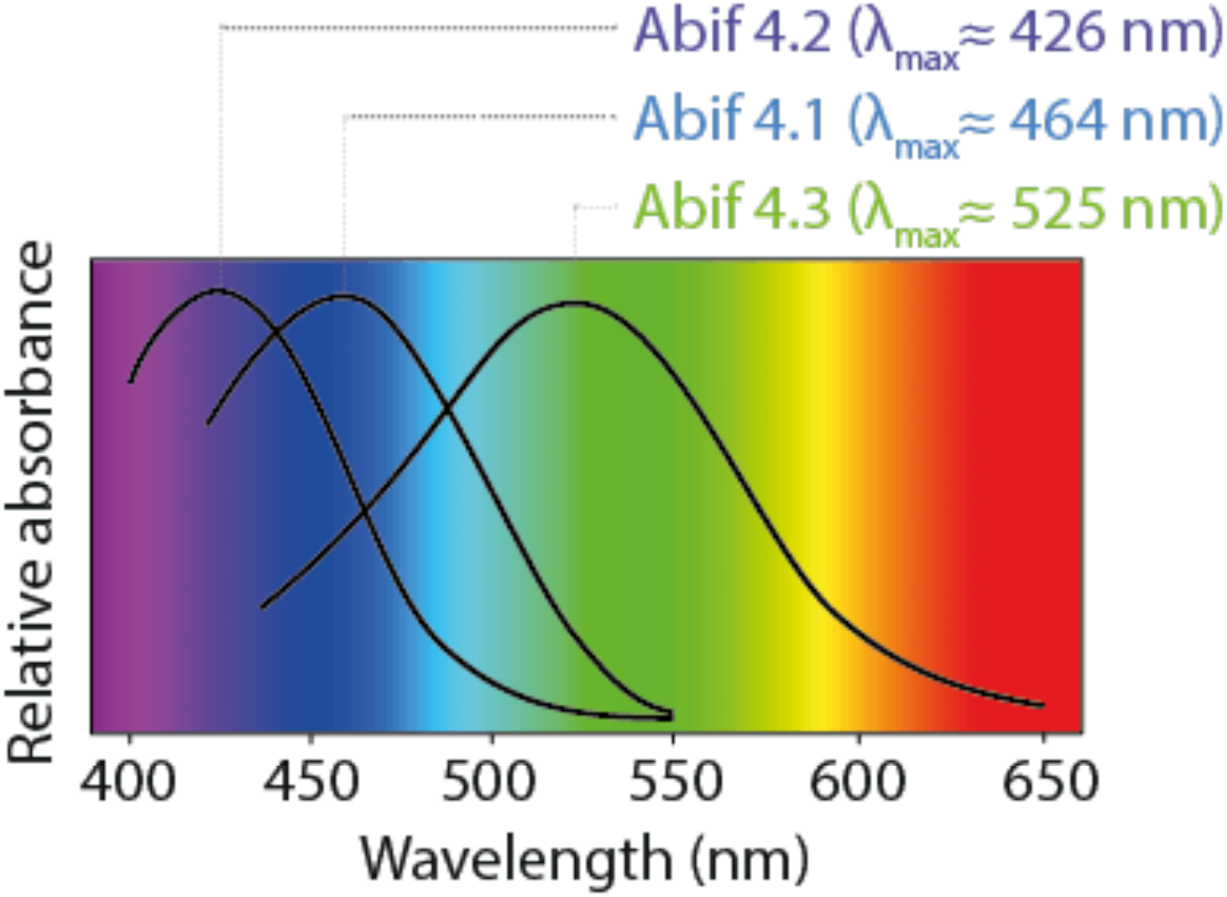
Schematic representation summarising the absorption spectra of the three opsins of *Antedon bifida* with the peak of light wavelength for the maximum absorbance.

### Potential photosensory structures in crinoids

Opsin immunostaining revealed the presence of at least two opsins in various tissue structures associated with the ambulacral grooves of the calyx and pinnules. One opsin was associated with the different nerve plexi whereas another opsin was expressed at the tips of the tube feet. These histological structures containing opsins are ideally located on the surface of the body and face away from the substratum, allowing a maximum detection of the sunlight rays.

The first opsin detected in the nervous plexus by antibodies raised against opsin 4 from the sea star *Asterias rubens* very likely corresponds to opsin 4.1 of *A. bifida*. Indeed, as evidenced by the alignments of the protein region recognised by the antibody, *A. bifida* opsin 4.1 shows the highest percentage (46.66%) of similarity with the epitope sequence of *Asterias rubens* rhabdomeric opsin (see the alignment in **Suppl. Fig. S2**). The presence of rhabdomeric opsins directly linked to the nervous system is relatively well described in other echinoderms, notably in the radial nerves and sensory nerves innervating the tube feet in sea urchins and brittle stars [Ullrich-Lüter et al 2011; Delroisse et al 2014]. It is well known that the ectoneural basiepithelial nerve plexus of echinoderms is involved in the transmission of sensory information [Weber and Grosmann 1977; García-Arrarás et al 2001; Mashanov et al 2023; Adameyko 2023], suggesting that this nervous plexus may also be involved in transmitting photo-sensorial information. By contrast, a photosensory activity within the hyponeural plexus is more intriguing. Indeed, this nervous plexus, located deeper in the connective tissue, is generally considered to perform only motor functions in echinoderms [García-Arrarás et al 2001]. Nevertheless, some paleontological studies have demonstrated that the hyponeural nerves in Paleozoic crinoids Camerata innervated directly some specific sensory organs in the calyx [Haugh 1978], demonstrating a possible dual sensory-motor role in crinoids, including a possible role for opsins in these deeper nerve fibres.

The detection of opsins in hyponeural nerves could shed new light on the still-unknown function of the saccules present below the surface of crinoid pinnules. These spherical bodies are highly refractive [Mallefet et al 2023] and are quite abundant in some species of feather stars, such as our study model *Antedon bifida*. They are regularly arranged under the epidermis on either side of the ambulacral grooves in the calyx and pinnules. The exact function of these saccules is still unknown, but numerous hypotheses about their physiological role have already been proposed, such as mucus secretion [Carpenter 1884; Clark 1921; Mironov and Pawson 2010], as an energy reserve [Hyman 1955] or playing a role in light production in bioluminescent species [Mallefet et al 2023]. Because of their optical properties, a hypothesis has been formulated suggesting their possible role as refractive lenses enabling light rays to converge on photoreceptors [Holland 1967]. The expression of rhabdomeric opsins in the hyponeural plexus located under the saccules, and innervating them, seems to support this hypothesis.

Earlier studies have also highlighted the presence of numerous pigments in the subepithelial connective tissue of *A. bifida* ambulacral grooves, as well as in other feather stars [Carpenter 1876, Dimelow 1958]. This pigmentation could potentially play a shading role at the level of ecto- and hyponeural nerve *plexi* expressing opsins. This shading system would be necessary to allow a directional photoreception, as demonstrated by phototactic tests in *A. bifida*.

The second opsin was specifically localised in the epidermis of the tube feet bordering the ambulacral grooves of the calyx and pinnules. This opsin was immunodetected using a different rabbit antibody raised against *Asterias rubens* opsin 1.1. Although this antibody recognises a ciliary opsin in the sea star, the epitope sequence shows a relatively high similarity with the opsin 4.2 from *A. bifida* (62.5%, **Suppl. Fig. S2**). Furthermore, a search for sequence homology has been performed against the entire *A. bifida* genome. Our results indicate that Abif-opsin 4.2 has a significant similarity (62.5%) with the sequence of the immunogenic peptide used to produce antibodies against sea star opsin 1.1. In the future, the results will need to be confirmed using immunoblots analyses.

Observing rhabdomeric opsin expression in tube feet of *A. bifida* is noteworthy, given that r-opsin localisation in tube feet has been previously demonstrated in other echinoderm classes such as sea urchins and brittle stars [Ullrich-Lüter et al 2011, Delroisse et al 2014], yet not in crinoids thus far. In contrast to other echinoderms, crinoids have their tube feet facing away from the substratum, which exposes them more effectively to light in the water column. Additionally, ciliary neurosensory cells have been found in various epidermal extensions such as “papillae” and “hillocks” of crinoid tube feet [Flammang and Jangoux 1991]. The opsin immunoreactivity is precisely located both at the base and occasionally at the tip of these sensory papillae, suggesting that these sensorial structures may be involved in light perception. This potential tube foot photoreception is corroborated by *pax 6* gene expression observed in the tube foot tips of the feather star species *Anneissia japonica* [Omori et al 2020], suggesting the probable presence of photoreceptors at the tips of tube feet in crinoids. Expression of *pax 6* is generally recognised for its pivotal role in the embryonic development of light-sensitive structures across bilaterian metazoans [Callaerts et al. 199f7; Gehring and Ikeo 1999; Arendt; 2003; Zuber et al. 2003; Martínez-Morales et al. 2004; Stierwald et al. 2004; Kozmik 2005]. Interestingly, *pax 6* has also been shown to be expressed in adult photoreceptor organs in various echinoderms, including co-expression of *pax 6* and opsin genes in the sea urchin species *S. purpuratus* [Ullrich-Lüter et al 2011]. However, it will be necessary to, e.g., express and characterise the function of these opsins by heterologous expression and develop gene knock-out workflows in this new model system to fully demonstrate the role and molecular mechanisms underlying non-canonical rhabdomeric opsins photoreception.

## Funding

YN and ED are supported by FRIA grants from the “Fonds de la Recherche Scientifique” of Belgium (F.R.S.-FNRS) and a PDR grant (T.0169.20). MAL is supported by the F.R.S-FNRS (MISU grant No F.6002.24). AL was supported by a FRIA grant from the F.R.S.-FNRS. PF is research director from the F.R.S.-FNRS. JD is a postdoctoral researcher funded by a PDR project from F.R.S.-FNRS (T.0071.23).

## Acknowledgements

This study is a contribution to the BioSciences Research Institute from the University of Mons and the “Centre Interuniversitaire de Biologie Marine” (CIBIM). The authors are grateful to the staff of the Marine Station of Concarneau for help and support.

## Conflict of interest

The authors declare no conflicts of interest.

## Author contributions

YN, IE, JD collected samples. YN, MAL, AL, LV and JD performed the experiments. MAL and YN performed *in silico* and *in vitro* opsin characterisation. YN, MAL and JD analysed and interpreted the results. YN and JD prepared the first draft of the manuscript. JD and PF supervised the study. All authors read and approved the final manuscript.

## Data availability

Data generated/analysed during the current study are available from the corresponding author upon reasonable request.

## Supplementary data

**Supplementary Table S1.** List of opsin sequences from bilaterian species used for the *in silico* and phylogenetic analysis.

**Supplementary Figure S1.** Phylogenetic analysis of crinoid opsins. (A) Global phylogenetic tree of the nine representative ancestral opsin lineages in bilaterian metazoans supported by maximum likelihood method. Bootstrap values are indicated at the root of each opsin cluster and represented further in the branch by a circle size proportional to the bootstrap value. Methodological details are presented in the legend of Figure 4.

**Supplementary Figure S2.** Multiple alignment, identity and similarity values between the two immunogenic peptides of the sea star (*Asterias rubens*) opsin 1.1 and opsin 4 recognised by antibodies and the crinoid (*Antedon bifida*) opsin sequences. Other opsin sequences of the sea star *Asterias rubens* and the sea urchin *Strongylocentrotus purpuratus* were added as model comparisons.

**Supplementary Figure S3.** Control tests for opsin immunolabelling in the ambulacral groove of the *A. bifida* calyx: (A) immunofluorescence detection of opsin 4.1 in the basiepithelial nerve plexus (opsin labelling in red and DAPI labelling of cell nuclei in blue). As saccules are highly refractive structures, they show strong red autofluorescence. (B) immunofluorescence detection of opsin 4.2 (in red) at the tip of tube feet. (C) Immunohistochemistry detection of opsin 4.2 (in brown) at the tip of tube feet. (D) Negative control in immunofluorescence without primary antibody. (E) Negative control in immunofluorescence using primary antibodies depleted with rabbit preimmune serum. (F) Negative control in immunohistochemistry without primary antibody.

## Notes

### Competing Interest Statement

The authors have declared no competing interest.

### Summary of Updates

Additional findings on the in vitro characterisation of Antedon opsins.

