## Supplementary figures and images for "Opsin-based photoreception in Crinoids: a molecular and behavioural study of *Antedon bifida*"

### Supplementary Figure S1

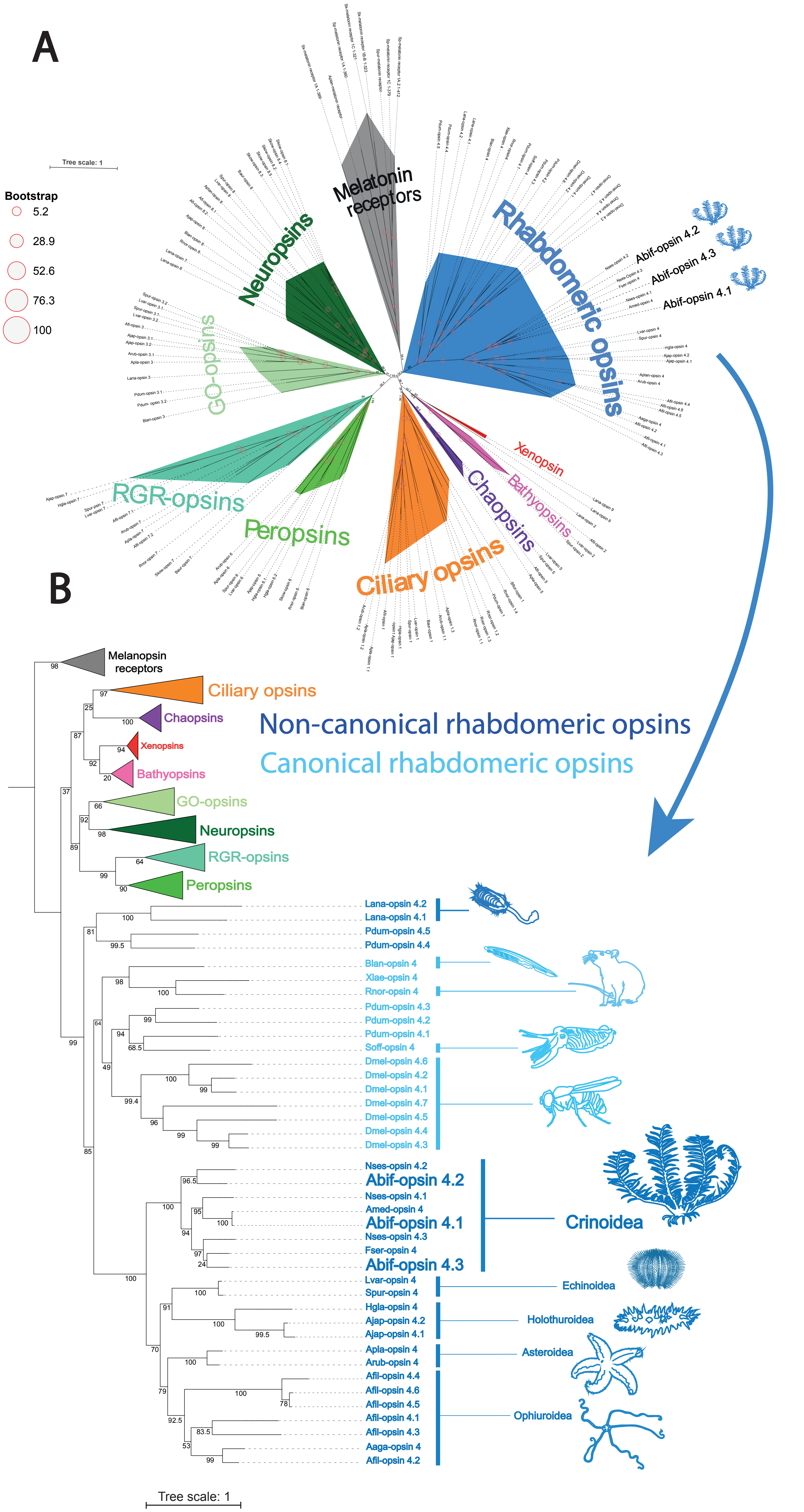

### Supplementary Figure S3

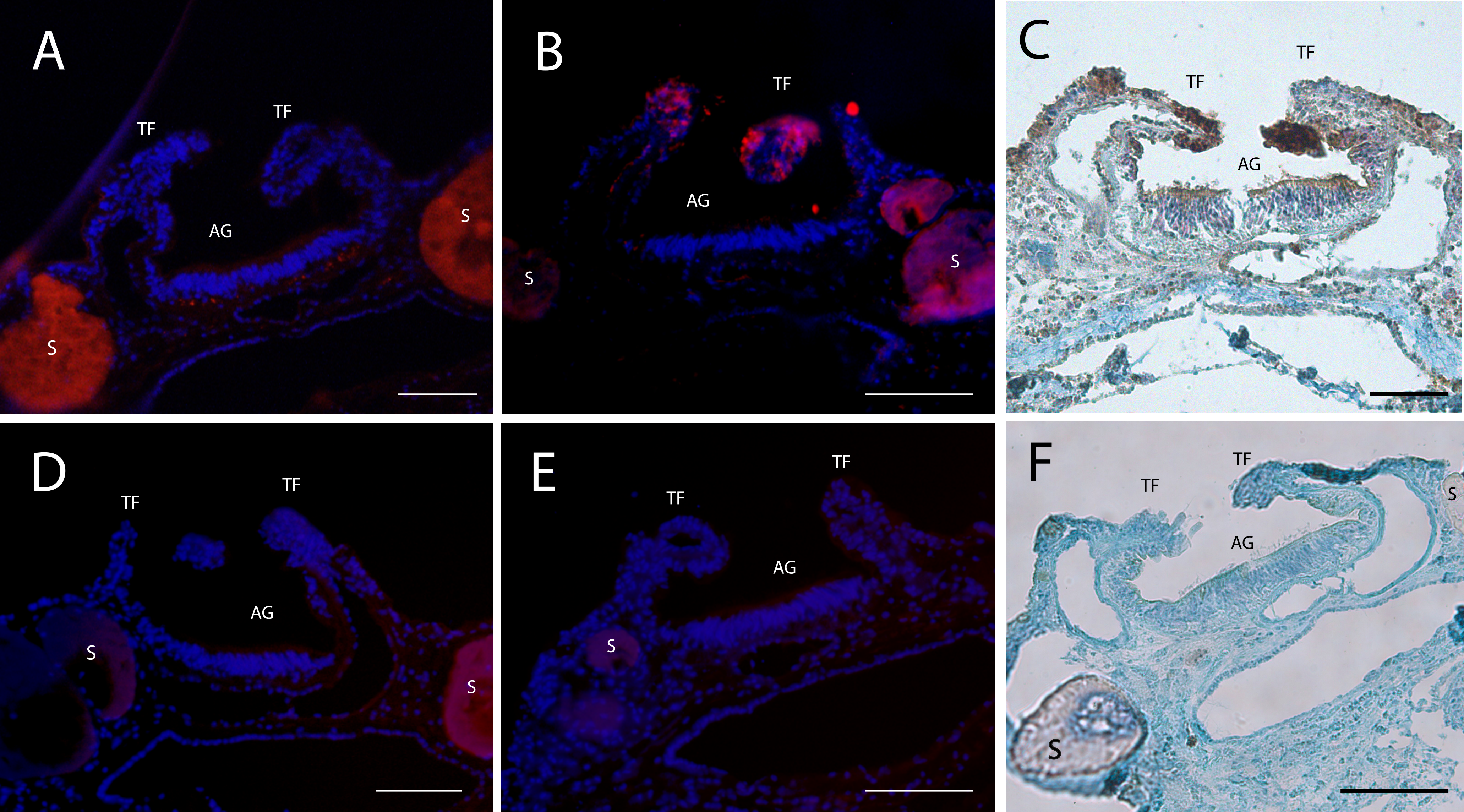
