## Supplementary Figure S2 for "Opsin-based photoreception in Crinoids: a molecular and behavioural study of *Antedon bifida*"

### Immunogenic peptide Arub-opsin 4

| | 450 … 460 | Identity | Similarity |
|---|---|---|---|
| *Antibody_peptid_opsin4_Arub* | ----NPAAVLNEKEELTKL |  |  |
| *Arub-opsin4* | AAYDNPAAVLNEKEELTKL | *100%* | *100%* |
| *Spur-opsin4* | ASASAPTPGVNDKEYLTKM | *46.66%* | *60%* |
| *Abif-opsin4.1* | GGIDNPAQLDN-----TDL | *40%* | *46.66%* |
| *Abif-opsin4.2* | ------------------- | *NA* | *NA* |
| *Abif-opsin4.3* | ----------------SDR | *NA* | *NA* |
| *Arub-opsin1.1* | ----------------SSL | *NA* | *NA* |

### Immunogenic peptide Arub-opsin 1.1

| | 160 … 170 | Identity | Similarity |
|---|---|---|---|
| *Antibody_peptid_opsin1.1_Arub* | ----AIGWSEFTVEGGGTSC---- |  |  |
| *Arub-opsin1.1* | VIPPAIGWSEFTVEGGGTSCSVNW | *100%* | *100%* |
| *Spur-opsin1* | CVPPLFGWNRYTYEGPGTACSVAW | *50%* | *56.25%* |
| *Abif-opsin4.1* | SVPPLFGFGRYTSEGYDLSCTFDY | *37.5%* | *50%* |
| *Abif-opsin4.2* | AVPPLLGFGRYTREGYGLSCTFDY | *43.75%* | *62.5%* |
| *Abif-opsin4.3* | SVPPLFGFGSYSREGYGLSCTFDY | *37.5%* | *56.25%* |
| *Arub-opsin4* | SLLPFFGLGAYVLEGYGVNCTFDY | *31.25%* | *43.75%* |
